# Finishing a complete giraffe genome from telomere to telomere with Verkko-Fillet

**DOI:** 10.1101/2025.10.01.679366

**Authors:** Juhyun Kim, Benjamin D Rosen, Sarah E Fumagalli, Kristen L Kuhn, Amy Long, Jeffrey J Schoenebeck, Heather Schwartz, Lan Wu-Cavener, Aleksey V Zimin, Douglas R. Cavener, Timothy P.L. Smith, Adam M. Phillippy, Sergey Koren, Arang Rhie

## Abstract

High-quality genome assemblies are essential for understanding speciation, evolution, and for building reference genomes and pan-genomes. Despite major advances in long-read sequencing technologies and graph-based assembly algorithms, current assemblies often remain incomplete and require refinement, including correction of haplotype switching and resolution of gaps, particularly within repetitive regions such as ribosomal DNA clusters and recent segmental duplications. These limitations largely stem from challenges in graph curation and are not adequately addressed by conventional polishing methods, which focus primarily on nucleotide-level corrections. To overcome these issues, we developed Verkko-Fillet, a Python-based interactive framework for genome graph inspection, editing, and curation that also provides step-by-step guidelines for downstream polishing. Verkko-Fillet takes as input Verkko output files, including the assembly graph, haplotype paths, Hi-C contacts, and additional ONT reads or alternative assemblies aligned to the graph, and offers tools for visualizing, modifying, and exporting curated assembly graphs. It enables users to track changes, resolve complex structural features, fill gaps, and enhance assembly quality beyond conventional polishing. As a case study, we applied Verkko-Fillet to the giraffe genome (*Giraffa tippelskirchi*), refining a draft assembly (QV 61.5) to a telomere-to-telomere (T2T) reference genome (QV 73.6) by improving both contiguity and completeness. Along with the T2T diploid giraffe genome, we provide gene annotations on this assembly as a valuable resource. Our results highlight the critical role of graph-based curation in producing high-quality, gapless, T2T assemblies that are ready for downstream biological analyses.

## INTRODUCTION

Generating high-quality genome assemblies is a critical foundation for understanding both individual species and their evolutionary relationships across the tree of life. A reference genome with minimal errors enables accurate functional annotation, comparative genomics, and evolutionary inference. To this end, numerous international efforts—such as the Telomere-to-Telomere (T2T) Consortium^1–3^, the Human Pangenome Reference Consortium (HPRC)^4,5^, Vertebrate Genomes Project (VGP)^6^ and Ruminant Telomere-to-Telomere (RT2T) Consortium^7^— focused on producing complete and accurate genome assemblies. These initiatives have driven the development of a wide range of genome assembler^8–11^ and polishing algorithms^12–14^ aimed at improving assembly continuity, correctness, and completeness.

Traditional *de novo* assembly methods relied on scaffolding assembled from long-reads into linear haploid sequences^15–19^. However, approaches have shifted toward graph-based assembly, which allows for more accurate representation of complex genomic regions by integrating linked-read, Hi-C or phasing information using trio^8,10,11,20–24^. These tools enable the accurate reconstruction of diploid^4,25–28^ and polyploid genomes^29–31^, and even precise haplotype phasing, avoiding the collapse of multiple haplotypes into a single consensus sequence—a limitation often inherent to linear assembly models.

Despite significant advancements in long-read sequencing technologies and genome assembly algorithms, persistent issues such as gaps and haplotype-switch errors remain in assembled genomes^32,33^. These problems are often caused by sequencing dropouts, long homozygous stretches between haplotypes, segmental duplications, and complex repetitive regions—particularly within rDNA clusters and centromeres^34,35^. While read mapping based polishing methods have been used in the past to fill or extend sequence into the gap^36–38^ and correct assembly errors^14,39,40^, they largely depend on the quality of the backbone assembly and are ineffective at resolving large, misassembled regions. Reads can be mis-mapped or prefer the higher quality sequence represented elsewhere, confounding the polishing results^6^. To maximize the recognized benefits of polishing, structural issues, common to Verkko’s initial, uncurated assembly output, are best addressed prior to mapping WGS reads. However, manually editing assembly graph paths to correct such regions—and subsequently generating the necessary files for re-building the assembly—is technically challenging and difficult to reproduce in a systematic way.

To address this challenge, we developed Verkko-Fillet, a Python-based interactive framework for genome assembly graph curation and gap resolution. Verkko-Fillet is designed to support multiple input formats commonly used in genome assembly workflows, including assembly graphs (GFA)^41^, ONT graph alignments (GAF)^42^, path files (GAF), and assembled sequences (FASTA). The core Verkko-Fillet object (hereafter vf-obj) integrates these inputs and maintains an exportable structured record of paths, nodes, edges, gaps, and modification history, allowing users to systematically track and manage changes throughout the curation process.

Furthermore, Verkko-Fillet provides compatibility with widely used visualization tools such as IGV^43^ and Bandage^44^, facilitating seamless transitions between analysis, visualization, and curation. It can also generate statistical graphics using its internal functions. Here, we present a complete “telomere-to-telomere” (T2T) reference genome of the giraffe (*Giraffa tippelskirchi*) constructed using Verkko-Fillet, demonstrating each step of the gap-filling and curation process in detail starting from a naive Verkko output. This assembly achieved substantial improvements in base-level accuracy (QV), genome completeness, and alignment quality, ultimately resulting in a gapless genome in which all contigs span from telomere to telomere. Comparative analysis with the previously published giraffe reference genome further supports the validity of our approach. The new assembly more accurately reflects known biological and cytogenetic features, including the orientation and composition of chromosomes as inferred from cattle karyotypes^45^. This underscores the utility of Verkko in combination with Verkko-Fillet for generating biologically accurate and complete reference genomes.

## RESULTS

### An interactive framework for generating T2T contigs

The Verkko assembler produces diverse data types across multiple layers, including assembly graphs, homopolymer-compressed nodes, stitched DNA segments (“pieces”), paths, and contigs (Figure S1). This complexity increases when assemblies are generated using Hi-C or trio-binning strategies. Effectively integrating these heterogeneous data types is essential for resolving gaps and achieving T2T contiguity, thereby enhancing overall assembly quality. Verkko-Fillet provides a unified framework to oversee the entire refinement process—from parsing the initial Verkko outputs to producing a gap-resolved assembly that is ready for polishing.

We developed the Verkko-Fillet object (hereafter referred to as “vf-obj”), a Python class designed to read and organize all output formats from Verkko into a single object using attribute-based methods (Figure S2), that seamlessly integrates all Verkko-generated data. The primary datasets handled by vf-obj are the assembly graph and paths. The graph consists of nodes and edges, while paths define the linear order and orientation of nodes corresponding to each contig. During gap-filling steps, vf-obj stores the name and orientation of nodes used to close assembly gaps and subsequently writes corrected path files for rerunning Verkko’s consensus module to generate an updated, gap-resolved assembly.

It is recommended to evaluate the quality of the assembly prior to path curation (Figure 1A). Verkko-Fillet provides functions to calculate QV by detecting error k-mers in the assembly, assess completeness by identifying telomeric sequences at contig ends, count remaining gaps, and assign chromosome names through alignment to a reference. It also analyzes the proportion of detectable telomeric motifs at the ends of contigs, as well as the number and location of gaps, to classify the assembly’s completeness. A chromosome is considered a “T2T contig” if both telomeres are present with no gaps, a “T2T scaffold” if both telomeres are present but one or more internal gaps remain, and just a “scaffold” if one or both telomeres are missing. These QC steps are recommended each time a new assembly is generated.

**Figure 1.**
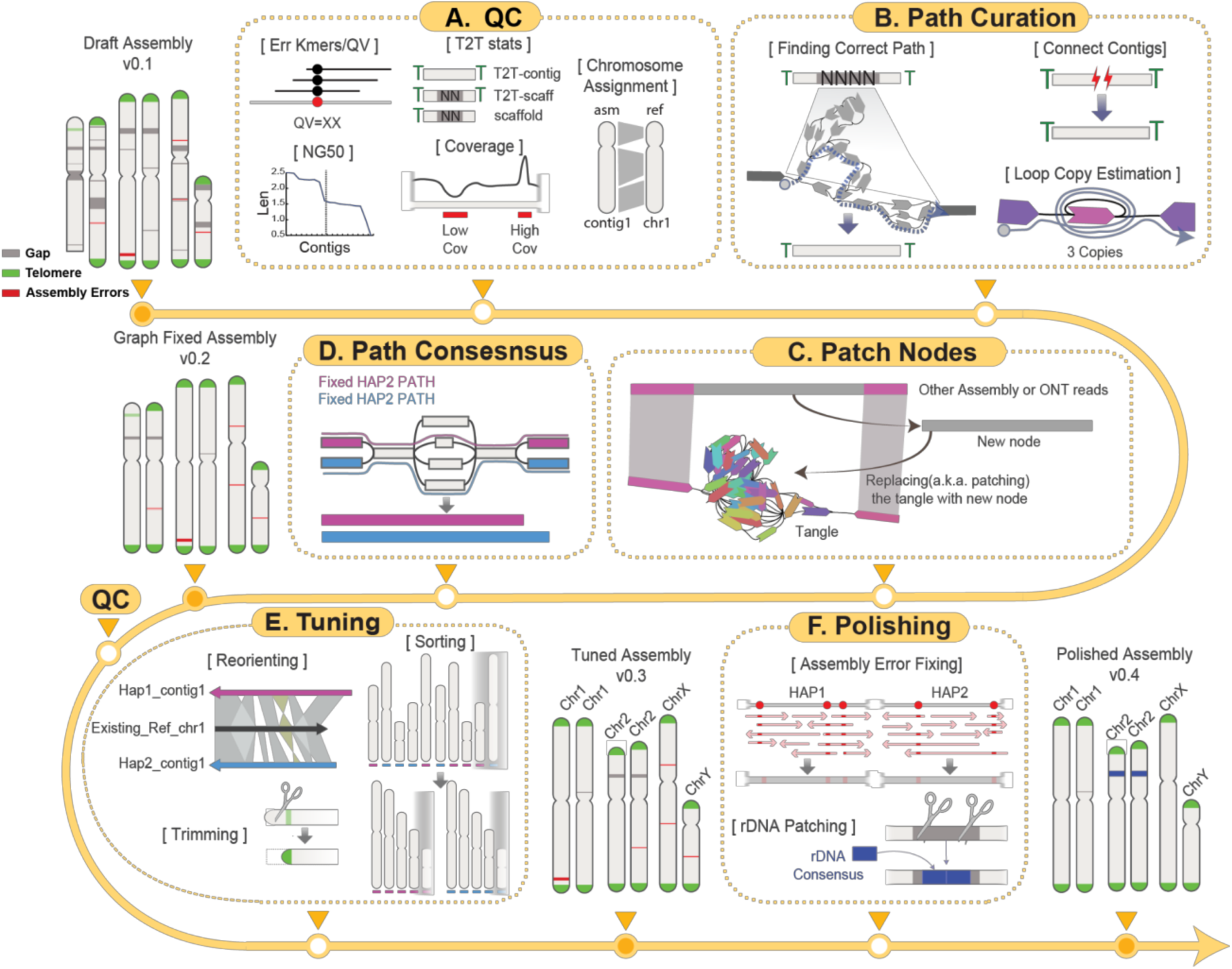
Verkko-Fillet is a software framework for the analysis and curation of complete genome assemblies. (A) The quality control (QC) step involves calculating error k-mers, assessing contig completeness, and performing chromosome assignment where applicable using a reference genome. (B) Gaps are resolved by tracing paths based on alignments of ONT reads to the assembly graph. Telomeric sequences at contig tips can be recovered by reconnecting broken paths into contiguous sequences. (C) New patching nodes are generated to restore intact paths by editing graph topology. Split reads and assemblies generated by other tools (e.g., Hifiasm^8^) are aligned to regions where the graph is broken between nodes, enabling the creation of new nodes for patching. (D) Verkko then generates a consensus assembly using the curated path files from steps (B) and (C), producing a gap-filled genome. (E) The tuning step applies contig-level modifications—including renaming, trimming, reorienting, and sorting—to generate a final, chromosome assigned T2T assembly. (F) The polishing step corrects base-level assembly errors using error k-mers and variant calling, while unresolved gaps in rDNA regions are addressed by rDNA patching, resulting in a finalized high-quality T2T assembly.

The core gap-filling function of Verkko-Fillet leverages ONT long-read alignments—specifically the ONT reads used in the initial assembly for scaffolding—mapped to the assembly graph with GraphAligner^46^. Reads spanning the flanking nodes of unresolved gaps are identified and used to infer the correct traversal paths through the graph, enabling replacement of missing regions (Figure 1B). This process incorporates copy number estimation of repeats within loops, reconnection of previously disconnected scaffolds, and construction of new bridging nodes from the extraction of sequencing information from multiple ONT reads (Figure 1C), which are subsequently used to regenerate consensus sequences (Figure 1D). Throughout the process, Verkko-Fillet supports exporting intermediate files in IGV- and Bandage^44^-compatible formats, facilitating simultaneous inspection of the assembly graph and sequence-level features.

The structurally refined assembly then undergoes tuning and polishing. The tuning step involves reorienting scaffolds according to known chromosomal orientation (if available), trimming, and sorting by chromosome or haplotype (Figure 1E). For species with no prior reference, Verkko-Fillet provides an option to sort the chromosomes by length and orient them based on user-provided centromere information. A finalized high-quality T2T assembly can then be generated by following the workflow provided in, a tutorial for base-level polishing using error k-mers and variant-calling methods (Figure 1F). rDNA regions are notoriously difficult to assemble due to their high copy number, so Verkko-Fillet includes a function to replace rDNA gaps with two copies of an rDNA consensus generated using Ribotin^47^, ensuring representation of these regions in the final assembly (Figure 1F).

Here, we present representative analyses performed at each step of the assembly refinement process using the genome of a male giraffe as a case study (Figure S3). The giraffe genome consists of 14 autosomes and the X and Y sex chromosomes, totaling 30 diploid chromosomes, along with mitochondrial DNA. Cytogenetically, all chromosome pairs except chromosomes 14 and Y are metacentric chromosomes, while chromosomes 14 and Y are acrocentric^48^. The initial draft assembly (T2T-mGirTip1v0.1) was generated using Verkko v2.2 with parental Illumina short reads to support trio-binning, followed by haplotype extension using Hi-C data. The initial assembly graph comprises 2,755 nodes and 3,679 edges, representing a total diploid genome size of 3.40 Gb. This graph-based assembly consists of 10 T2T-contigs, 17 T2T-scaffolds, and 3 scaffolds classified as non-T2T (Figure 2A, Figure S4).

**Figure 2.**
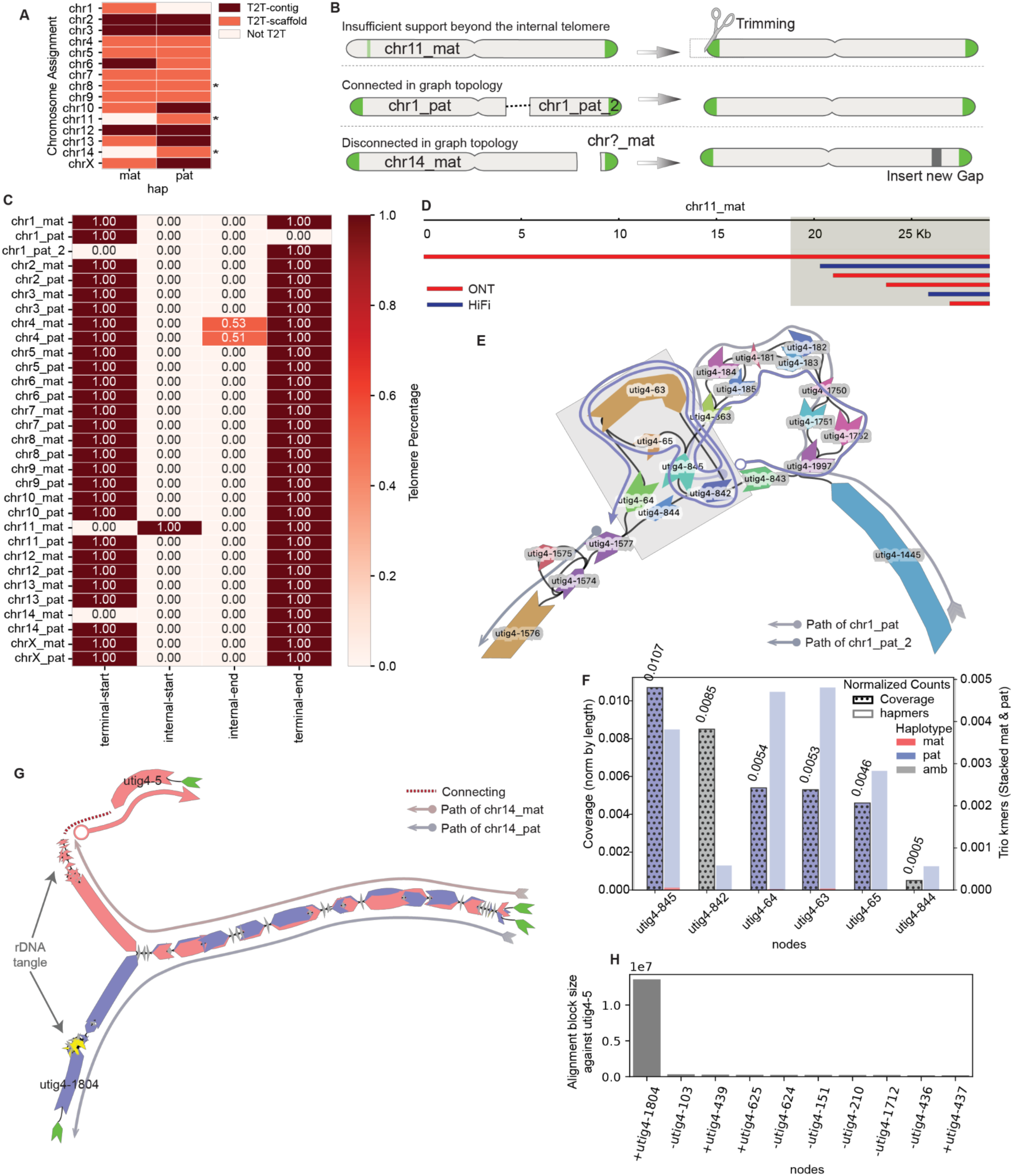
Recovering telomere-to-telomere (T2T) contigs. (A) Heatmap of T2T status for each chromosome in the T2T-mGirTip1v0.1 assembly. Scaffolds are classified as T2T contigs (red), T2T scaffolds (orange), or non–T2T (beige). Asterisks (*) denote contigs containing rDNA arrays. (B) Schematic representation of three categories of non–T2T contigs: (1) insufficient support beyond an internal telomere, (2) a contig fragmented into two scaffolds despite being connected in the graph topology, and (3) a scaffold missing terminal nodes containing telomeres, even when represented in the graph topology. (C) Maximum proportion of telomeric sequence per scaffold, distinguishing between terminal ends (≤15,000 bp from the chromosome end) and internal regions (>15,000 bp from the end). (D) Sequencing read composition at the tip node of maternal chr11 in the T2T-mGirTip1v0.1 assembly, with reads colored by sequencing technology: HiFi (blue) and ONT (red). The yellow shaded box highlights regions containing canonical telomeric repeats. Only a single ONT read supports the non-telomeric sequence, likely representing a misplaced read with minimal support. (E) Assembly graph near a break in the paternal chr1 scaffold. Semi-transparent arrows indicate paths assigned to chr1 (paternal haplotype) in the T2T-mGirTip1v0.1 assembly: one from *utig4-1445(+)* to *utig4-842(-)*, and another beginning at *utig4-1577(-)*. The solid colored arrow indicates the final incorporated path. (F) Normalized coverage and hampers (haplotype-specific k-mer) counts per node length surrounding the breakpoint in panel E, highlighted by the gray box. (G) Graph structure of chr14 from both haplotypes. Nodes are colored by haplotype (maternal, red; paternal, purple). Telomeric nodes are marked with green triangles, and rDNA repeat regions are highlighted with arrows. The red dotted line indicates a novel connection between two contigs, while the solid pink arrow shows the newly incorporated path. (H) Pairwise alignment of the disconnected node *utig4-5* in panel G against all other nodes in the graph, ranked by relative alignment block length, to identify its homologous counterpart on the alternate haplotype.

### Recovering missing telomere ends

There are three common scenarios in which a contig fails to meet the criteria for classification as T2T (Figure 2B). First, misplaced reads at the contig tip may introduce additional non-telomeric sequence. Second, a contig path may become disrupted within a complex tangle—often caused by repetitive regions—resulting in its fragmentation into two separate scaffolds, even though they remain connected in the graph topology. Third, a contig may lack terminal nodes, leaving the ends unconnected in the graph topology. All three scenarios are amenable to correction through tuning and graph-based refinement, enabling recovery of complete T2T contigs.

We detected an anomalous pattern in the maternal contig of chromosome(chr) 11 (Figure 2C) when analyzing the distribution of telomeric motifs across contig termini. A high density of telomeric repeats was observed ∼18.8 kb from the 5′ end of the contig, while the true terminal sequence showed no detectable telomeric signal. Examination of read support at the 5′ terminus revealed that this non-telomeric segment was derived from a single ONT read, whereas the internal region enriched for telomeric repeats was consistently supported by both ONT and HiFi datasets (Figure 2D). We therefore inferred that the additional sequence at the contig tip originated from a misplaced ONT read. To recover a true T2T chromosome, the contig was trimmed upstream of the validated telomeric signal during the tuning step, after generating the gap-fixed consensus, because the reads could not be re-placed in node space (Figure 1E, Figures 1S).

We identified two large paternal haplotype contigs, both assigned to chr1, with one containing the p-arm telomere and the other the q-arm telomere (Figure 2C, E). These contigs should form a single continuous sequence but were separated by a large, complex tangle, presenting two possible traversal paths: utig4-64 or utig4-844. To resolve this ambiguity, we evaluated trio-specific k-mer counts and ONT read coverage for each node within the tangle and found that utig4-64 had a higher number of paternal k-mers and greater ONT coverage than utig4-844, indicating it more likely represents the true haplotype path (Figure 2F). Based on this evidence, we first identified the node within the large loop and then inferred the most plausible traversal as [utig4-842 → utig4-63 → utig4-64 → utig4-1577], thereby connecting the two contigs into a continuous paternal chr1 (chr1_pat) scaffold.

Maternal chr14 (Chr14_mat) was also missing one telomere (Figure 2C). Unlike the chr1_pat example, the chr14_mat graph lacked the telomeric node beyond the rDNA tangle (Figure 2G). However, we identified utig4-5, an unconnected node with a length comparable to utig4-1804—the telomere-bearing node at the end of the paternal short arm. To determine whether utig4-5 belonged to maternal chr14, we aligned it against all other nodes in the graph and ranked the results by alignment block length. The top match was utig4-1804, supporting the hypothesis that utig4-5 represents the missing maternal telomeric segment (Figure 2F). Because no edges connected utig4-5 to the main contig, we bridged the inferred path with a gap placeholder, yielding the structure: [utig4-5 → GAP → utig4-436]. With this adjustment, the contig now qualifies as a T2T scaffold.

### Gap filling

Gaps in genome assemblies can arise from multiple sources and can be broadly classified into two major categories (Figure S5): graph tangles and unassigned haplotypes, excluding the “missing telomeres” described earlier. Tangles occur when nodes in the assembly graph are interconnected in a highly complex manner, making it difficult to resolve a clear, haplotype-specific path. These typically occur in repetitive regions of the genome such as recently duplicated gene families or satellite repeat arrays.

Gaps in the giraffe genome were resolved by integrating multiple sources of supporting data. The primary dataset consists of ONT long-read alignments to the assembly graph (Figure 3A). Specifically, we re-aligned the ONT reads used in the initial assembly to the graph using GraphAligner^46^, enabling the identification of reads that span the flanking nodes on either side of a gap, as well as node-level coverage profiles. These spanning reads provide strong evidence for the correct traversal through unresolved regions. Additional data sources, including Hi-C read alignments and trio-specific k-mer markers, were also employed when available to further support gap resolution and facilitate accurate haplotype assignment.

**Figure 3.**
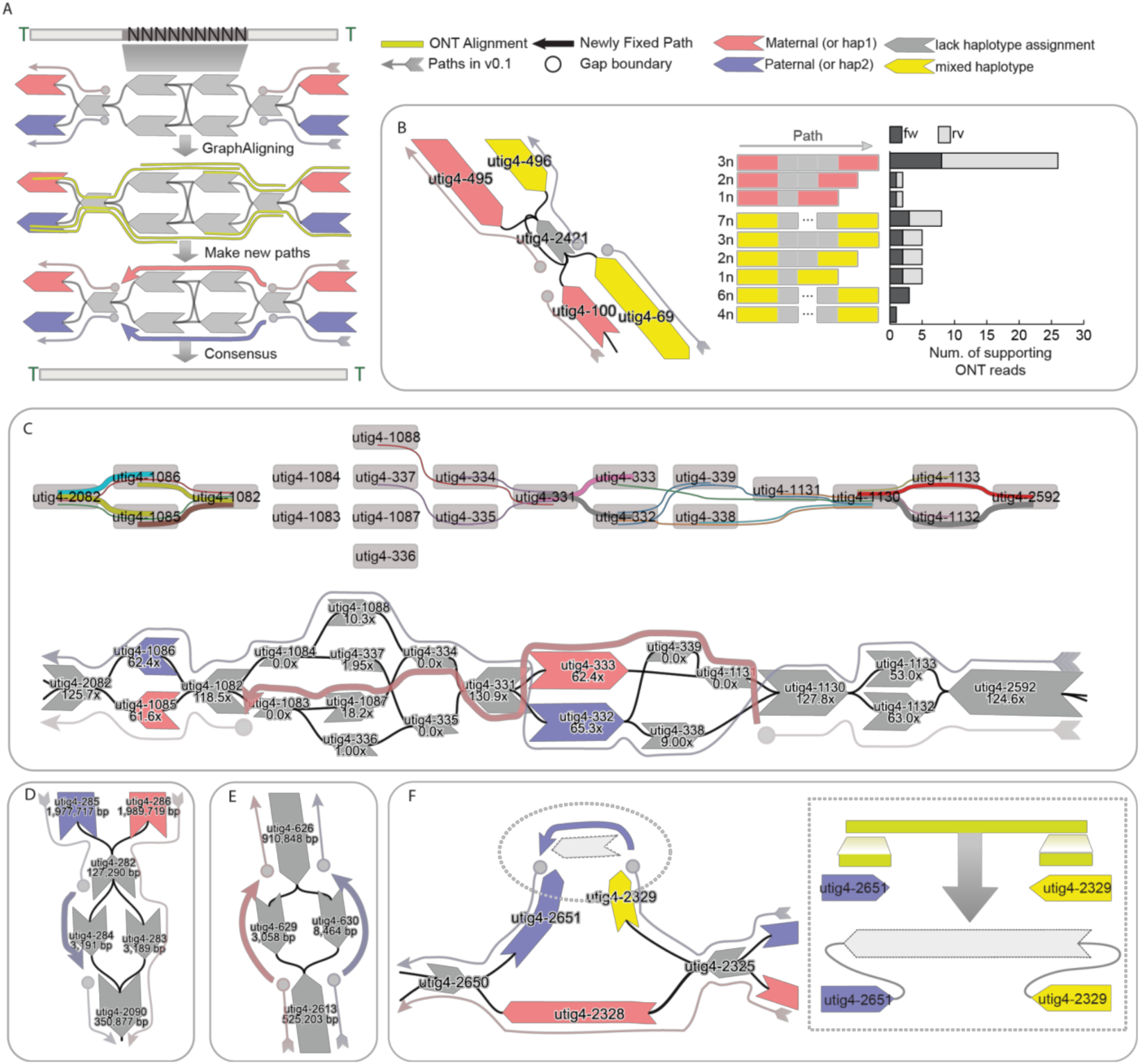
Generating a gapless assembly. (A) Schematic of gap filling using ONT alignments on the assembly graph. Colors indicate haplotypes: maternal (pink), paternal (purple), mixed trio markers (yellow), and nodes lacking trio k-mers (gray), which may represent homozygous nodes or repeats. Gaps arise when a contig path cannot form a continuous linear sequence. The key idea is to reconstruct an intact path by leveraging ONT alignments or assemblies from other tools mapped onto the assembly graph. (B) Example of copy number estimation for a repeat node. The stacked bar chart shows the number of ONT reads supporting flanking haplotype-specific nodes containing the repeat, stratified by alignment strand. (C) Example of a complex tangle in the graph, where ONT alignments span multiple nodes. (D) Example of haplotype assignment by inference from the path of the counterpart haplotype. (E) Example of ambiguous cases where haplotypes are assigned randomly to both haplotypes. (F) Patching example. A missing connection between two nodes on the paternal haplotype was resolved by generating a new patch node derived from split ONT alignments.

Loop resolution was straightforward when sufficiently long ONT reads span the haplotype-specific flanking nodes surrounding a loop, supporting estimation of the number of repeat units traversed by each read (Figure 3B). In the maternal haplotype, 25 ONT reads were found to span from utig4-495 to utig4-100, each traversing three copies of utig4-2421. In contrast, for the paternal haplotype, 8 supporting reads spanned utig4-69 to utig4-496, each traversing seven copies of utig4-2421. Based on these estimates, we reconstructed the loop by inserting the corresponding number of repeat nodes into each haplotype-specific path.

We resolved the path in more complex tangles by integrating coverage and ONT alignment information. In this case, the diploid graph diverged into four nodes within the same genomic region (Figure 3C). ONT alignments were detected only once each on utig4-1088 and utig4-334, respectively, but HiFi coverage indicated substantially higher support for utig4-1088 and utig4-1087, both of which were ultimately included in the resolved paths. Consequently, utig4-1087 was incorporated into the maternal path to fully resolve the tangle, since utig4-1088 had already been assigned to the paternal haplotype.

Each path in the assembly graph is highly specific to its corresponding haplotype. Consequently, gaps may arise when there is more than one node that is not homologous and not assigned to a haplotype in the path (Figure 3D, E). In such cases, the gap can often be resolved by leveraging the corresponding path from the alternate haplotype (Figure 3D).

However, when the gap is due to uncertainty or ambiguity in both haplotypes, as a last resort it may be necessary to fill the gap by assigning a path randomly, which could achieve a more contiguous, complete sequence albeit at the expense of possibly introducing a haplotype switch error (Figure 3E).

Gaps may also arise from missing nodes or edges in the assembly graph, often due to insufficient HiFi coverage or lack of support from ONT long reads (Figure 3F). In this case, the graph was disconnected, which prevented ONT alignments from spanning both nodes in a single path. However, we identified ONT reads that aligned uniquely to both nodes in a split manner, allowing us to reconstruct the missing sequence. This process required introducing a new node and edges to connect the flanking nodes, thereby restoring graph continuity, and the new node is then used during the consensus step to fill the gap.

In summary, T2T-mGirTip1v0.1 contained 284 initial paths. Using Verkko-Fillet gap filling, the T2T-mGirTip1v0.2 assembly retained only 31 essential paths—28 autosomes, two sex chromosomes, and one mitochondrial chromosome. Of the 45 gaps identified in the initial assembly, we successfully filled all but one, introducing a single new gap to reconstruct a fully telomere-capped scaffold for chr14_mat (Figure 2G).

### Trimming, chromosome assignment and polishing

Consensus sequence generation (Figure 1D) for T2T-mGirTip1v0.2 was followed by tuning steps including trimming, reorientation (flipping), renaming, and sorting (Figure 1E). chr11_mat was the only contig missing one telomeric end (Figure S6). As noted above, the chr11_mat contig required specific trimming during this tuning stage because the tip sequence was derived from the consensus of a single ONT read (Figure 2C).

Each contig was renamed according to its chromosomal assignment and haplotype, based on alignments to a matched reference genome when available. Using the available giraffe reference (ASM1759144v1), we assigned contigs to chromosomes based on sufficiently long alignment blocks (Figure 4A). However, approximately one-third of the sequences could not be reliably oriented because the alignments contained mixed directionalities (Figure 4A). We leveraged the conventionally used comparative karyotype of giraffe to cattle^45,49^ to resolve orientation by aligning the cattle reference genome (Bos taurus, GCF_002263795.3) to T2T-mGirTip1v0.2 (Figure 4B). For example, the known cattle–giraffe homology for giraffe chr1 is BTA6(–) followed by BTA8(+), which is consistent with the alignment of T2T-mGirTip1v0.2. In contrast, cattle BTA1, BTA28, and BTA26 aligned in the opposite order on T2T-mGirTip1v0.2 compared with the established recombination karyotype, prompting us to flip this contig. Overall, 9 of the 15 contigs per haplotype required reorientation to conform to the widely accepted giraffe karyotype. Finally, the contigs were sorted by chromosome name and haplotype.

**Figure 4.**
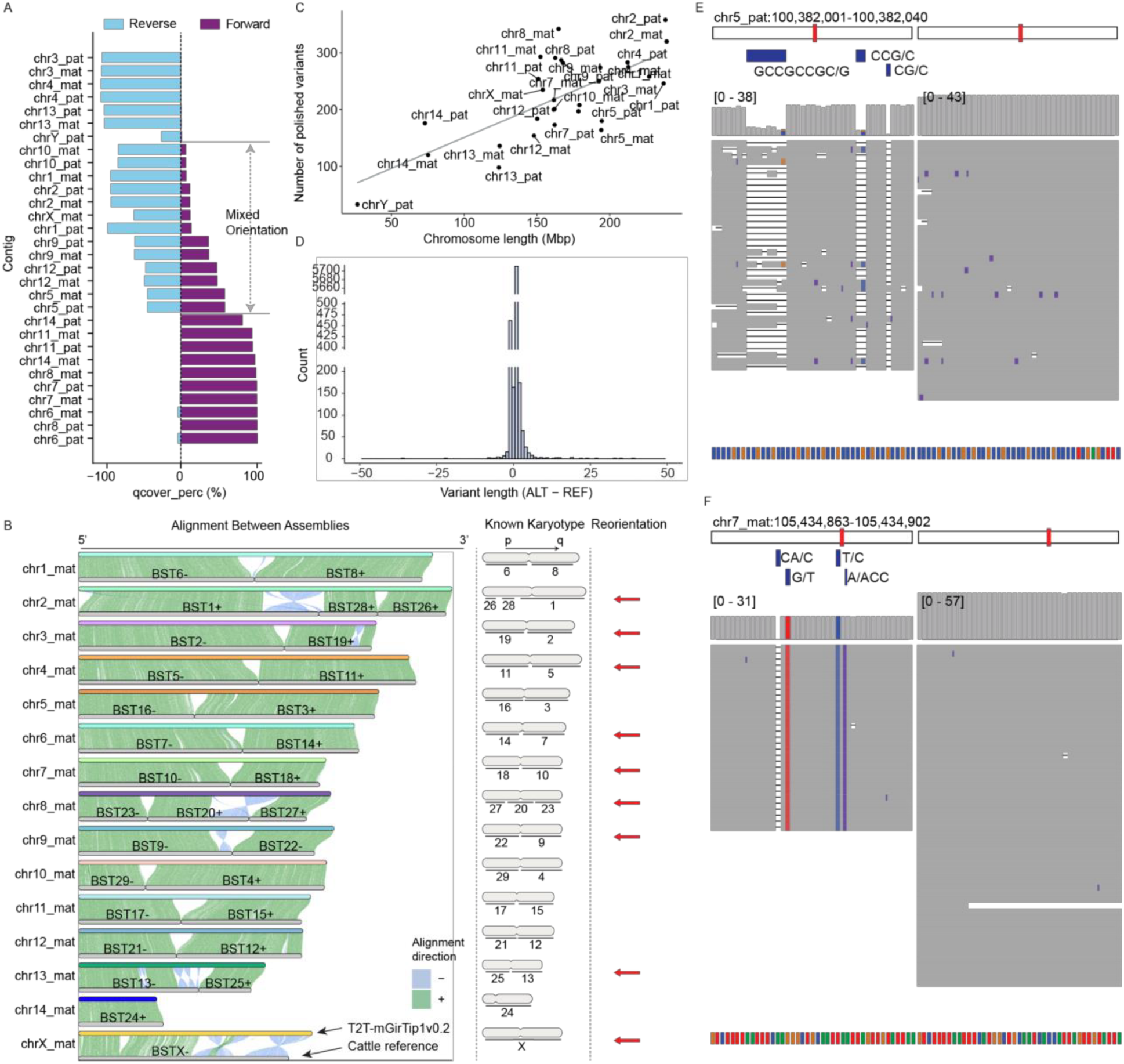
Tuning and Polishing of the T2T-mGirTip1v0.2 assembly. (A) Alignment of the query assembly (T2T-mGirTip1v0.2) to the species-matched reference (ASM1759144v1) for chromosome assignment and orientation checking. This step identifies which contig corresponds to each chromosome and whether it should be flipped. Several contigs show mixed-strand alignments within a single chromosome, creating ambiguity relative to the existing reference. (B) Alignment of the cattle reference (ARS-UCD2.0) to T2T-mGirTip1v0.2 to ensure contig orientations are consistent with the known giraffe karyotype based on cattle homology. The upper panel shows T2T-mGirTip1v0.2-Mat contigs, and the lower panel shows cattle chromosomes. Contigs requiring reorientation are marked with red arrows in the “Reorientation” column. (C) Number of polished variants per chromosome as a function of contig length. The regression line is shown as y = 42.1 + 1.09X; R² = 0.51. (D) Histogram of polished variants by size. Variants of size 0 correspond to single-nucleotide substitutions, values >0 represent insertions, and values <0 represent deletions. (E, F) Examples of the same genomic position before polishing (left) and after polishing (right). HiFi read alignments are shown to illustrate mismatches, indels, and coverage. Base-resolution information is provided at the bottom: A (green), C (blue), G (orange), and T (red).

Base-level correction of the assembly (Figure 1F) was performed using a variant-based strategy. ONT reads were aligned to the diploid genome, while hybrid alignments of HiFi and Illumina reads were performed against each haploid assembly (maternal and paternal) as well as diploid assemblies. Variants were then called and filtered to identify sequence discrepancies unsupported by the underlying read data, following the approaches described by McCartney et al.^14^. and Yoo et al.^50^. In total, 6,689 corrections were identified (excluding those on chrM), corresponding to an average of 1.3 variants per Mb (Figure 4C). The majority of corrections were +1 indels (Figure 4D). Polishing improved local read alignments, reducing mismatches and indels within corrected regions (Figures 4E, F).

The current constraints of ultralong read length and accuracy limit the ability to resolve complex rDNA arrays for some species’ assemblies. Verkko-Fillet provides an option to replace rDNA gaps with a model rDNA sequence (morph), in the form of [gap – rDNA units – gap], thereby incorporating rDNA sequences into the assembly (Figure S7). This conservative approach enhances the assembly by minimally preserving chromosome-specific rDNA unit information while avoiding introduction of assembly errors created by inaccurate node traversals. In T2T-mGirTip1v0.2, all gaps, including those within rDNA tangles, were resolved without the need for this replacement. However, one rDNA unit on chr14_pat still exhibited a high density of local error k-mers even after polishing, indicative of an error in the node that was chosen and was thus removed (Figure S8).

Finally, the mitochondrial genome was circularized, and pseudoautosomal regions (PARs) were identified on chrX and chrY. All these post-polishing steps were carried out using Verkko-Fillet modules.

### Evaluating Assembly quality following structural correction and base-level polishing

The structural improvements made with Verkko-Fillet were assessed for the T2T-mGirTip1v0.3 genome. Coverage profiles for both HiFi and ONT reads were broadly consistent, with expected low coverage restricted to telomeres or satellite enriched regions with known sequencing biases (Figure 5A). Notably, in T2T-mGirTip1v0.1, Flagger^4^ statistics and alignment patterns revealed that not only near the gap but also ∼200 kb of adjacent regions exhibited alignment difficulties (Figure 5B). After gap filling, read alignments improved substantially. Haplotype-specific alignments (MAPQ > 0) were restored at the boundary of node utig4-2295, and coverage profiles stabilized across both HiFi and ONT datasets. Importantly, no residual anomalies were detected by either Flagger or the coverage issue track (https://github.com/arangrhie/T2T-Polish/blob/master/coverage/), which reports abnormal coverage based on dinucleotide composition and error k-mers.

**Figure 5.**
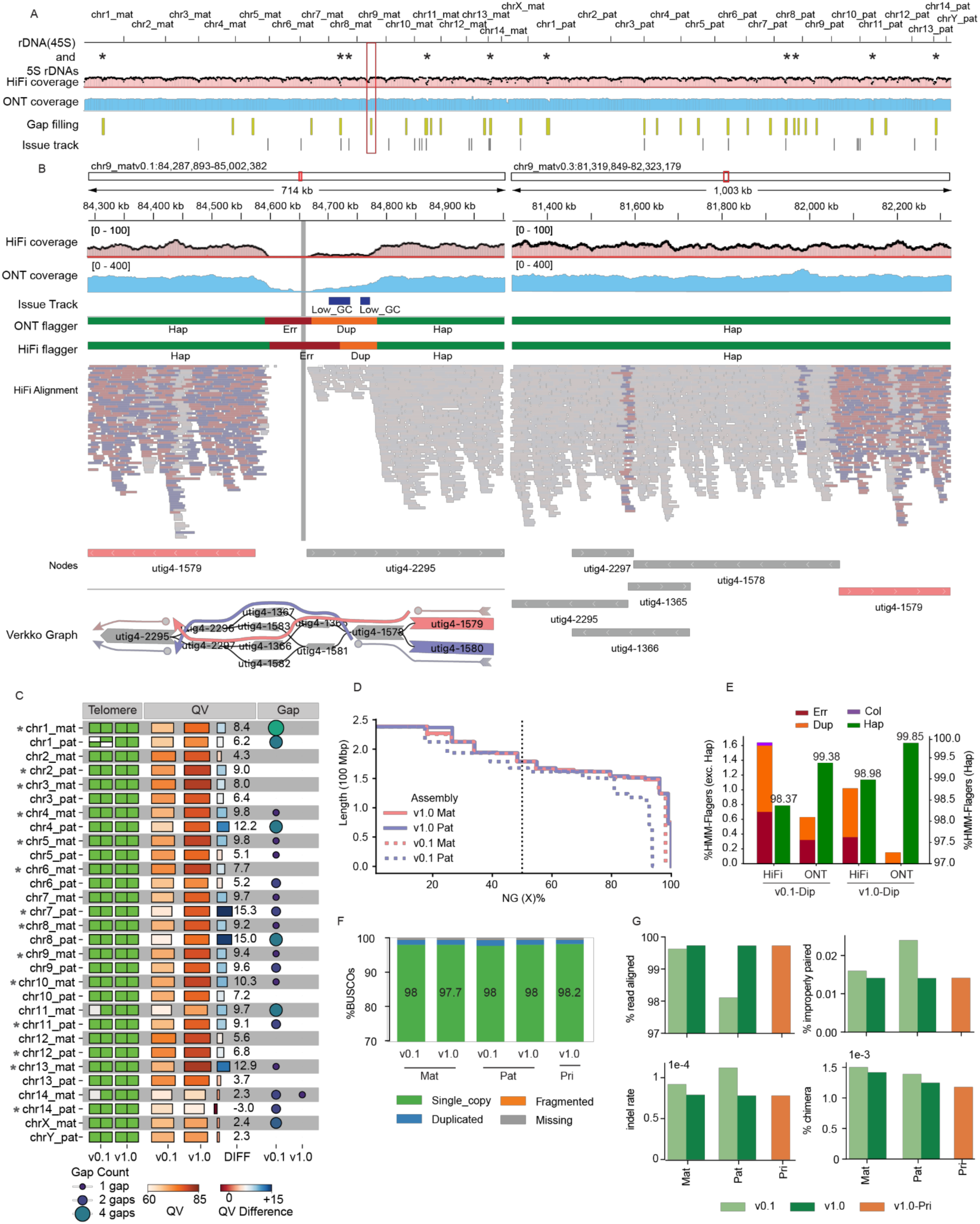
Structural and Base-Level Refinement of the T2T-mGirTip1 Assembly Using Verkko-Fillet. (A) Overview of the T2T-mGirTip1-v0.3 structurally corrected genome after gap filling and tuning. HiFi and ONT alignments show genome-wide stable coverage from both platforms, except at rDNA loci, centromeres and telomeres. The HiFi coverage track is overlaid with the first allele (red dots) and second allele (black dots) from NucFlag^69^. Gap-filled regions are shown on the “Gap filling” track. The “Issue” track highlights regions identified by error k-mers and abnormal HiFi/ONT coverage. (B) Zoomed-in genomic plot around a gap-filling event on chr9_mat, highlighted by the red box in panel A. Comparisons are shown for T2T-mGirTip1-v0.1 (left) and T2T-mGirTip1-v0.3 (right). Coverage abnormalities classified by hmm-flagger are shown, with HiFi read alignments colored by strand. Shaded colors indicate reads mapped with MAPQ = 0. The node sequence alignment with the original graph and reconstructed paths is shown at the bottom. The original gap is highlighted in gray in T2T-mGirTip1-v0.1. Note that chr9 was among the contigs flipped during the tuning step (Figure 3B). (C) Summary metrics comparing T2T-mGirTip1-v0.1 and T2T-mGirTip1-v1.0, including T2T statistics, QV values, number of gaps, and QV improvements. Columns: (1, 2) telomere detection on p- and q-arms; (3, 4) QV values for each assembly; (5) QV improvement; (6, 7) number of gaps. Primary contigs are denoted with an asterisk (*) before the contig name. (D) NG(X)% by version and haplotype. Dotted lines represent v0.1; solid lines represent v1.0. The total length of v1.0 for each haplotype was used as the genome size for the corresponding haplotype assembly. (E) Percentage of coverage abnormality categories from flagger by version and read type. Reads were aligned to the diploid genome. (F) Stacked bar plot of BUSCO^51^ gene estimation by version and haplotype (including the primary assembly). The percentage of “Single copy” genes is highlighted in text on the bar plot. (G) Statistics of Illumina short-read alignment by version and haplotype (https://broadinstitute.github.io/picard/).

The overall impact of all changes made from the initial (v0.1) to the final polished (v1.0) assembly were assessed by comparing various assembly quality metrics (Figure 5C, Table S1). In v1.0, all chromosomes contained telomeres at both ends, with only a single remaining gap on chr14_mat, introduced during contig joining. The overall QV also improved substantially, increasing from an average of 61.5 to 73.6. The largest improvement was observed on chr7_pat, where the QV increased from 62.0 to 77.4 (Figure 5C). The haploid, linear representation of the genome to be used as a reference was constructed by selecting a primary haplotype for each chromosome based on having fewer gaps and higher QV, yielding assemblies hereafter referred to as T2T-mGirTip1v1.0-Pri.

Assembly contiguity improved, with N50 values increasing from 148 Mb to 162 Mb for the maternal haplotype and from 150 Mb to 162 Mb for the paternal haplotype (Figure 5D, Table S2). Flagger^4^ statistics showed incremental improvements from v0.1 to v1.0 with lower collapses and duplications in both HiFi and ONT reads (Figure 5E). The v1.0 assembly exhibited fewer errors, collapsed regions, and duplications, while also showing a higher proportion of haploid-specific regions (Figure 5E, Table S2). Gene content assessed with BUSCO^51^ results were in line with Flagger, such that v1.0-Pri contained the highest proportion of single-copy gene predictions (98.2%), along with fewer missing or incomplete genes (%) (Figures 5F, Figure S9). Finally, mapping of Illumina short-read data from the mGirTip1 to each assembly version further demonstrated v1.0 yields higher alignment rates, fewer improper pairs, reduced indel rates, and fewer chimeric alignments (Figure 5H). Overall, as graph refinement, tuning, and polishing progress, the assembly quality is substantially improved, making it more useful for downstream analyses by enhancing gene content and mapping quality.

### The T2T-mGirTip1v1.0 genome provides a more accurate and biologically valid representation of giraffe chromosomes

We next compared T2T-mGirTip1v1.0-Pri to the latest publicly available giraffe reference genome, derived from a male Rothschild’s giraffe ASM1759144v1 (*Giraffa camelopardalis rothschildi*)^52^, assembled with ONT reads and scaffolded with Hi-C (Figure 6A). Compared with ASM1759144v1, the T2T-mGirTip1v1.0-Pri assembly demonstrates notable quality improvements (Table 1).

**Figure 6.**
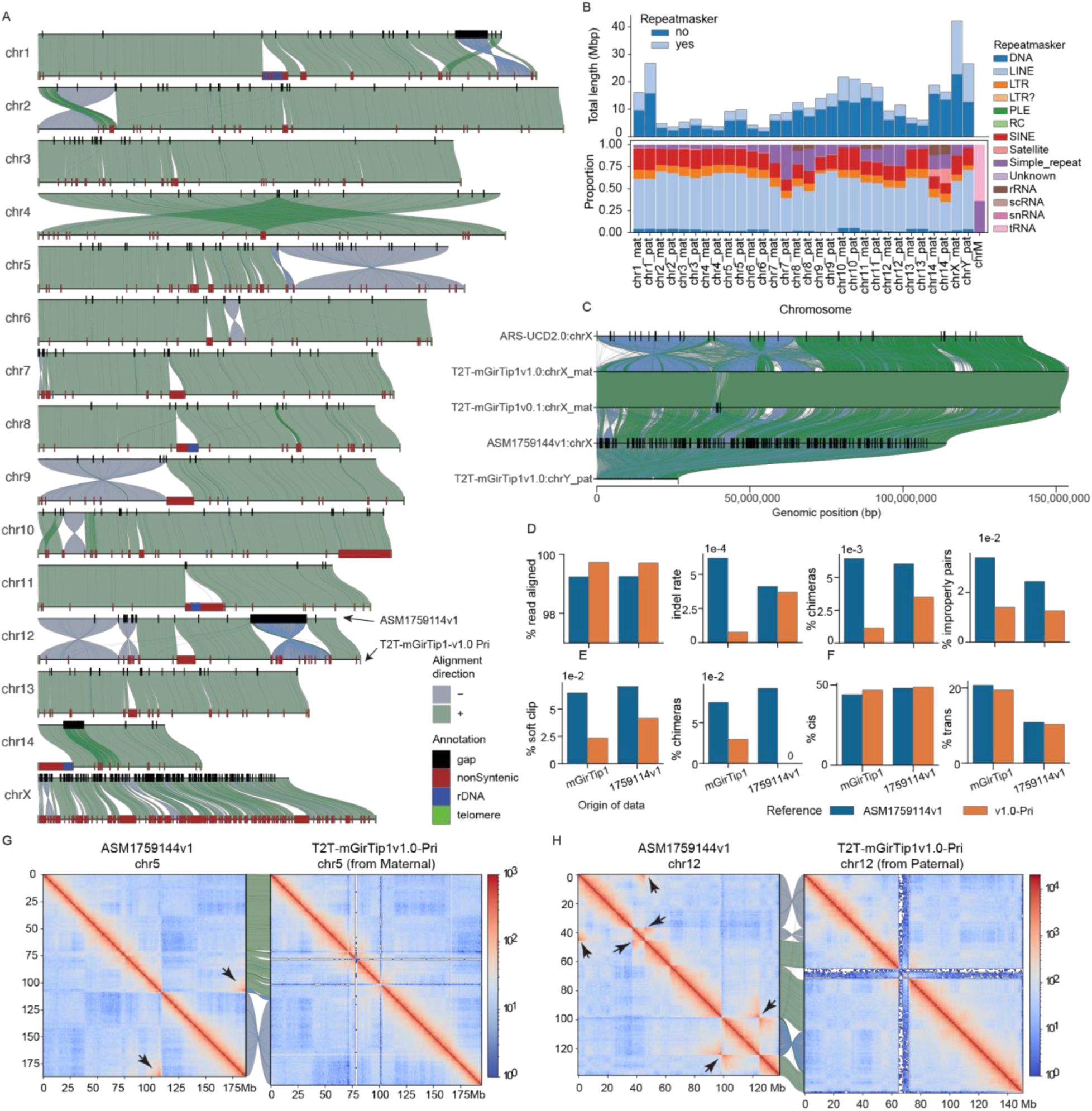
From draft to complete: the T2T-mGirTip1-v1.0 giraffe genome assembly. (A) Syntenic plot showing structural differences between the T2T-mGirTip1-v1.0 primary assembly and the giraffe reference genome ASM1759114v1, generated using SVByEye^73^. Top: ASM1759114v1 with gaps shown in black boxes. Bottom: T2T-mGirTip1-v1.0 primary contigs, with non-syntenic regions (brown), rDNA (blue), and telomeres (green). (B) Total length of non-syntenic regions per chromosome (upper panel), with darker colors indicating repeat-covered regions identified by RepeatMasker^72^. The lower panel shows the proportion of each repeat type within the non-syntenic regions for each contig. (C) Syntenic comparison of the chrX from cattle (ARS-UCD2.0) and giraffe (T2T-mGirTip1-v1.0 and ASM1759114v1), and of the chrY from T2T-mGirTip1-v1.0. Gaps (black) and telomeric regions (green) are marked on each contig. (D) Direct and cross-validation statistics of Illumina short-read alignment on the haploid genomes T2T-mGirTip1-v1.0-pri and ASM1759114v1. The x-axis indicates the dataset origin (mGirTip1 or ASM1759114v1). (E) Comparison of ONT alignment statistics between assemblies. (F) Hi-C sequence data from each dataset aligned to the haploid genome, showing the percentage of cis- and trans-interactions. Cis refers to read pairs separated within chromosomes, and trans refers to read pairs spanning between chromosomes^70^. (G, H) Hi-C contact maps aligned to ASM1759114v1 and T2T-mGirTip1-v1.0 for chr5 (G) and chr12 (H). The long range interactions were highlighted with black arrows.

**Table 1.**
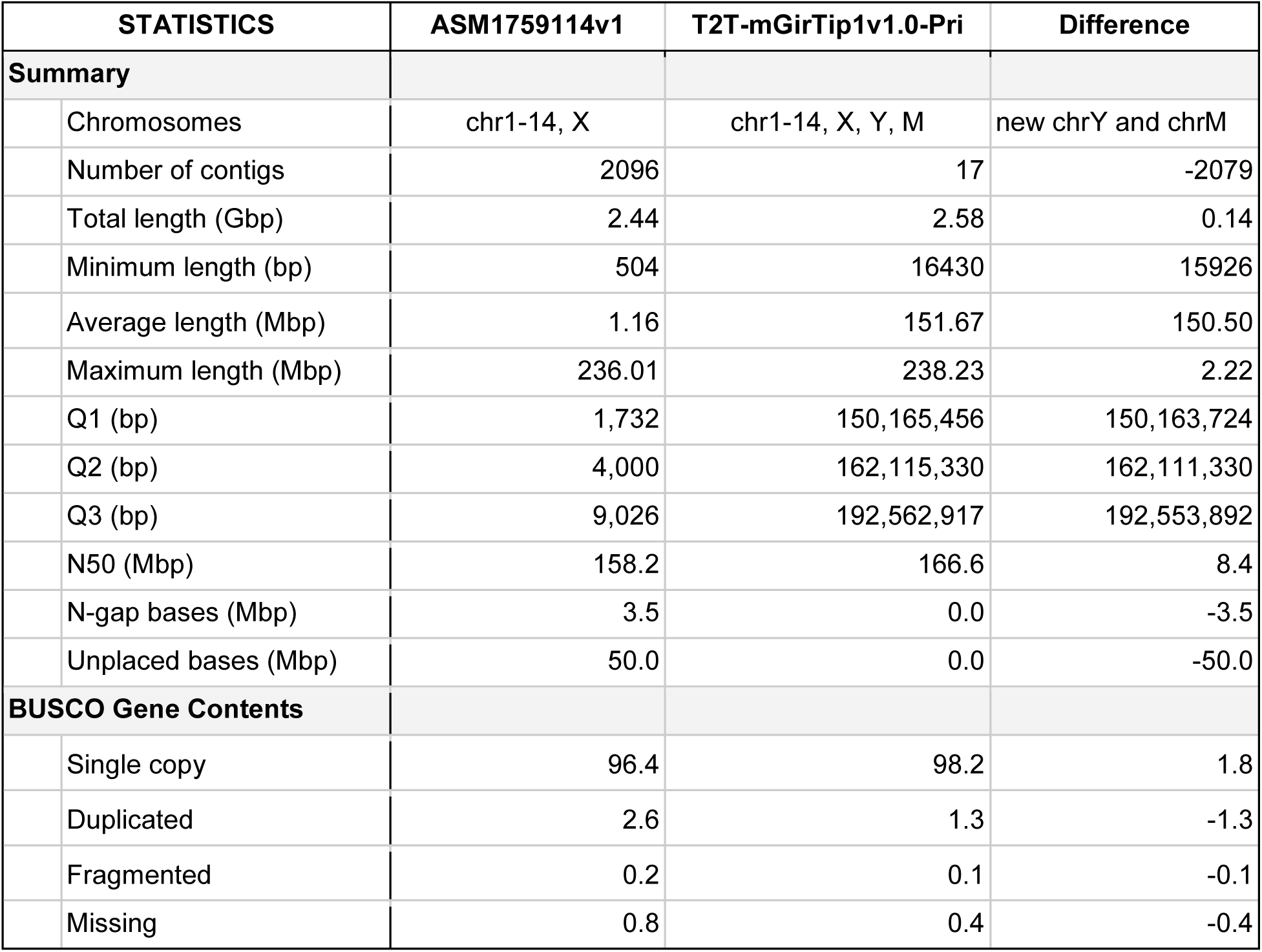
Comparison of ASM1759114v1 and T2T-mGirTip1v1.0-Pri giraffe genome assemblies.

| STATISTICS |  | ASM1759114v1 | T2T-mGirTip1v1.0-Pri | Difference |
| --- | --- | --- | --- | --- |
| **Summary** |  |  |  |  |
|  | Chromosomes | chr1-14, X | chr1-14, X, Y, M | new chrY and chrM |
|  | Number of contigs | 2096 | 17 | -2079 |
|  | Total length (Gbp) | 2.44 | 2.58 | 0.14 |
|  | Minimum length (bp) | 504 | 16430 | 15926 |
|  | Average length (Mbp) | 1.16 | 151.67 | 150.50 |
|  | Maximum length (Mbp) | 236.01 | 238.23 | 2.22 |
|  | Q1 (bp) | 1,732 | 150,165,456 | 150,163,724 |
|  | Q2 (bp) | 4,000 | 162,115,330 | 162,111,330 |
|  | Q3 (bp) | 9,026 | 192,562,917 | 192,553,892 |
|  | N50 (Mbp) | 158.2 | 166.6 | 8.4 |
|  | N-gap bases (Mbp) | 3.5 | 0.0 | -3.5 |
|  | Unplaced bases (Mbp) | 50.0 | 0.0 | -50.0 |
| **BUSCO Gene Contents** |  |  |  |  |
|  | Single copy | 96.4 | 98.2 | 1.8 |
|  | Duplicated | 2.6 | 1.3 | -1.3 |
|  | Fragmented | 0.2 | 0.1 | -0.1 |
|  | Missing | 0.8 | 0.4 | -0.4 |

Whole-genome alignment between T2T-mGirTip1v1.0-Pri and ASM1759144v1 revealed 220 Mbp of non-syntenic regions, of which 36.8% were composed of repeat sequences (Figure 6A, B; Table S3; Figure S10). The overall repeat composition across chromosomes was largely similar, with LINEs and SINEs comprising the majority of repetitive elements (Figure 6B). However, certain chromosomal regions displayed notable enrichment, including rRNA repeats on chromosomes 8, 11, and 14, and tRNA repeats on chr14 (Figure 6A, B).

Most notably, representation of the sex chromosomes was substantially improved in the T2T-mGirTip1v1.0 assembly (Figure 6C). The previous reference genome, ASM1759144v1, completely lacks the chrY and contains 288 gaps scattered across the chrX, totaling 145 kbp out of 114 Mbp. These gaps are likely the result of collapsed sequences from both sex chromosomes into the chrX, which in turn contributed to the numerous gaps and fragmented syntenic regions observed between chrX and chrY^53,54^. In contrast, T2T-mGirTip1v1.0 fully resolves both the chrX and chrY, without any gaps, with newly added bases of 42.21 Mb (27.4%) and 26.62 Mb (100%), respectively. Comparative alignment with the cattle chrX confirmed two previously reported inversions on the p arm of the giraffe chrX^45,55^.

We aligned Illumina, ONT and Hi-C data from both mGirTip1 and ASM1759144v1 to both assemblies (Figure 6D, E). Alignment of Illumina data showed that T2T-mGirTip1v1.0 had a higher proportion of properly aligned reads, along with lower proportions of chimeric reads, indel rates, and improper pairs compared with ASM1759144v1 (Figure 6D). Similarly, ONT data alignment revealed fewer reads with soft clipping and chimeric signatures when mapped to T2T-mGirTip1v1.0 (Figure 6E). Hi-C alignment further demonstrated improved performance, with T2T-mGirTip1v1.0 showing a higher number of cis interactions and fewer trans interactions than ASM1759144v1 (Figure 6F).

Extensive synteny was observed between the T2T-mGirTip1v1.0 assembly and ASM1759144v1 across many chromosomes, yet several chromosomes exhibited notable orientation discrepancies within specific regions (Figure 5A, Figure S11–S12). In particular, chr1, 2, 5, 6, 9, 10, and 12 displayed large-scale structural differences, including apparent inversion and translocation errors (Figure 6A, G–H; Figure S13). For example, the Hi-C contact map of chr5 in ASM1759144v1 revealed a large inversion on the q-arm. In contrast, chr12 in ASM1759144v1 contained multiple inversions, whereas no long-range interaction signals indicative of an inversion was detected in the T2T-mGirTip1v0.1 contact map (Figure 6H, Figure S14). Notably, Hi-C data from ASM1759144v1 aligned to ASM1759144v1 reproduced these long-range interactions, whereas the same signals were absent when aligned to T2T-mGirTip1v1.0, supporting the structural accuracy of T2T-mGirTip1v1.0 and suggesting that T2T-mGirTip1v1.0 represents a more reliable reference for the giraffe species (Figure S13, S14).

Gene annotation on the T2T-mGirTip1v1.0 assembly was performed with EviAnn^56^. Illumina RNA-seq and PacBio Kinnex read data were used as transcriptomic evidence, and a set of 576,821 mammalian proteins served as protein homology evidence from UniProt-SwissProt database^57^. We identified 20,848 protein-coding genes and 2,785 non-coding genes (Table 2). In total, 85,186 isoforms were predicted as protein coding, of which 83,131 were functionally annotated and 61,891 showed strong homology to known proteins in the primary assembly. BUSCO^51^ analysis derived from this new annotation further confirmed that T2T-mGirTip1v1.0 is highly complete in terms of gene content, based on the mammalia database (cetartiodactyla_odb10) (Figure S4, Table S4).

**Table 2.** Annotation results for primary and alternative haplotype assemblies of the T2T-mGirTip1v1.0 genome.

|  |  | T2T-mGirTip1v1.0 |  |
| --- | --- | --- | --- |
| Statistics |  | Primary (Autosomes + XY and MT) | Alternative |
| **Genes** |  |  |  |
|  | Protein-coding | 20,848 | 19,824 |
|  | Non-coding | 2,785 | 2,817 |
|  | Processed pseudogenes | 409 | 346 |
| **Transcripts** |  |  |  |
|  | Protein coding | 85,186 | 82,481 |
|  | Functionally annotated | 83,131 | 80,465 |
|  | Long non-coding RNA | 2,877 | 2,911 |
|  | Processed pseudo | 417 | 351 |
| **Proteins** |  |  |  |
|  | Unique proteins | 61,891 | 59,786 |

Collectively, T2T-mGirTip1v1.0 represents the first high-quality T2T giraffe genome, correcting many large inversion errors present in previous references and adding new sequences that were previously difficult to assemble using the Verkko and Verkko-Fillet workflows. We also provide both gene and transcriptome annotations for the diploid assembly, offering a valuable resource for biological research.

## DISCUSSION

Multiple genome assembly tools, such as Verkko^10,11^, Hifiasm^8^, and others, employ graph-based representations, where nodes denote sequences, edges represent their connectivity, and paths define haplotype-resolved contigs. Despite the advantages of these frameworks, manual curation of assembly graphs remains limited. As a result, unresolved gaps and structural errors frequently persist into final consensus sequences, requiring correction prior to polishing to avoid mismapping of the reads.

This limitation is addressed by Verkko-Fillet, a guided Python-based interactive tool that integrates seamlessly into the genome assembly workflow and can be easily installed via pip or in a conda environment. Verkko-Fillet provides graph-based quality control, manual path curation, error correction, and export of refined graph structures, thereby bridging the gap between the Verkko assembler and downstream polishing steps to generate high-quality, contiguous assemblies. Our results show that substantial improvements in QV can be achieved through graph correction and gap filling, even prior to polishing (Table S1). Consistently, alignment scores and coverage uniformity across sequencing platforms—including Illumina, HiFi, and ONT—were markedly improved. These findings demonstrate that resolving gaps and correcting structural errors in the assembly graph before polishing significantly enhances overall assembly quality. In addition, Verkko-Fillet supports post–gap-filling, tuning, and polishing workflows, including chromosome renaming, reorientation, rDNA patching, base-level error correction, and other procedures required to produce a finalized, complete assembly.

The input data representation in Verkko-Fillet is the Verkko genome graph, which consists of nodes and edges, along with associated paths that represent assembled sequences. Users can edit these paths to correct assemblies, guided by supporting evidence such as haplotype information, coverage profiles, and alignments from Hi-C and ONT reads. While this approach could in principle be extended to other assemblers that employ graph-based representations, it is currently limited to the Verkko framework. Support for other assemblers could be a potential area of future development.

We demonstrate the complete workflow for generating the telomere-to-telomere giraffe genome (T2T-mGirTip1v1.0) using Verkko-Fillet, highlighting improvements in QV, gene content, and NG50, supported by detailed examples. Comparative analyses showed that T2T-mGirTip1v1.0 preserves the expected DNA homology with the cattle genome and revealed several large inversion errors in the existing giraffe reference. In addition, we identified 220 Mbp of new sequences non-syntenic to the prior best assembly of a giraffe species, including proximal centromeric and rDNA sequences, as well as intact T2T contigs for chrX, chrY, and chrM, which may provide new insights into speciation. Beyond the high-quality assembly, we also provide a preliminary gene annotation for the diploid genome, including 40,672 protein-coding genes. As no RefSeq annotation of the Giraffe species is currently available in NCBI, this resource will be of substantial value for studies of giraffe conservation and ruminant biology.

## RESOURCE AVAILABILITY

### Lead contact

For additional information, resources, requests should be directed to the lead contact, Arang Rhie, Ph. D

### Materials availability

The giraffe samples are available upon request.

### Data and code availability

● Verkko-Fillet is a pip-installable Python package, publicly available at https://github.com/jjuhyunkim/verkko-fillet along with a full documentation.
● Sequencing data is available under BioProject PRJNA1367195, including samples from blood (SAMN51204129) and testes (SAMN51283588) from mGirTip1, as well as maternal (mGirTip2), paternal (mGirTip3) and nephew of mGirTip1 (mGirTip4) samples (SAMN51204130, SAMN51204131, and SAMN53299408 respectively).
● The T2T-mGirTip1v1.0-Mat, Pat, Pri, and Alt assemblies have been deposited under BioProject PRJNA1322564 and will be publicly available soon. In the meantime, the assemblies are available to download on genomeark (https://www.genomeark.org/t2t-all/Giraffa_tippelskirchi.html).
● The genome assembly and gene annotation data are available through the project’s GitHub repository (https://github.com/jjuhyunkim/giraffeT2T), which includes known issues and annotation resources.

## ACKNOWLEDGEMENTS

We thank the Cincinnati Zoo and the Nashville Zoo for providing samples for the giraffe family. We thank members of the Ruminant T2T consortium (https://www.ruminant2t.org) for conceptualizing and providing insightful comments during the development of this work. J.J.S. was supported by the Biotechnology and Biological Sciences Research Council (BBSRC) BBS/E/D/10002070. L.W.C. and D.R.C. was supported by Eberly College of Science and the Huck Institute of the Life Sciences, Penn State University. A.V.Z. was supported by the National Science Foundation (NSF) IOS-2432298. This work utilized the computational resources of the NIH HPC Biowulf cluster (https://hpc.nih.gov). This research was supported in part by the Intramural Research Program of the National Institutes of Health (NIH). The contributions of the NIH authors are considered Works of the United States Government. The findings and conclusions presented in this paper are those of the authors and do not necessarily reflect the views of the NIH or the U.S. Department of Health and Human Services. The results reported here were made possible with computational resources provided by the USDA-ARS SCINet initiative. Any mention of trade names or commercial products is solely for the purpose of providing specific information and does not imply recommendation or endorsement by the U.S. Department of Agriculture. The USDA is an equal opportunity provider and employer.

## AUTHOR CONTRIBUTIONS

J.K. designed Verkko-Fillet and performed all data analyses and drafted the manuscript. B.D.R. generated the initial T2T-mGirTip1v0.1 Verkko assembly. J.K., B.D.R., S.E.F., and J.J.S. optimized Verkko-Fillet. A.L and H.S. collected samples. D.R.C., L.W.C., and K.L.K. generated sequencing data. A.V.Z. conducted gene annotation. S.K. supervised compatibility between Verkko-Fillet and Verkko. A.R. provided feedback on the polishing step and handled data submission. T.P.L.S., A.M.P., S.K., and A.R. supervised the project. J.K. and A.R edited the manuscript with assistance of all authors. All authors read and approved the final manuscript.

## METHODS

Detailed methods are provided in the online version of this paper and include the following:

● KEY RESOURCES TABLE
● EXPERIMENTAL MODEL AND SUBJECT DETAILS

○ Sample collection
● METHOD DETAILS

○ Data generation
○ T2T-mGirTip1v0.1 assembly
○ Verkko-Fillet Functions and graph curation
○ Run Verkko consensus using the final path with filling the gaps
○ Tuning and polishing
● QUANTIFICATION AND STATISTICAL ANALYSIS

○ Assembly assessment
○ Genome annotation

## SUPPLEMENTAL INFORMATION

## METHODS

### EXPERIMENTAL MODEL AND SUBJECT DETAILS

#### Sample collection

Blood and tissue samples were collected from a Masai giraffe (*Giraffa tippelskirchi)* family at the Cincinnati Zoo and Botanical Garden, Cincinnati, OH USA and Nashville Zoo, Nashville, TN USA. For the paternal genome (Kimba, mGirTip3; DOB 2007), the skin samples were collected post-mortem from his legs that were saved and frozen (-4℉) and gDNA was isolated. He died in 2019 after an anesthetic procedure to assess chronic lameness issues. He was 12yrs old at the time of death. He was overall healthy other than his hoof issues causing the lameness. For the maternal sample (Tessa, mGirTip2; DOB 2006), tail hair root cells were collected when the animal was 15 years old and gDNA isolated. For the offspring (Fennesey, mGirTip1; DOB 2019), testes sample was collected when the animal was 2 years old during the castration procedure, and flash frozen in liquid nitrogen to preserve RNA integrity. Later, ∼12 cc of EDTA whole blood was collected from the left jugular vein with a 21 ga butterfly catheter and vacutainer setup when mGirTip1 turned 5 years old, and immediately frozen in liquid nitrogen to preserve chromosomal DNA integrity. Tessa and Fenn are both at the Cincinnati Zoo and are healthy. An older sibling of mGirTip1 (Nasha; DOB 2014), who is sharing the same parents as mGirTip1, was shipped to the Nashville Zoo in 2015. She gave birth to a calf in 2021, mGirTip4, who died from a neck injury along the cervical vertebrae shortly after birth. Postmortem tissues were collected from the cardiac muscle that were saved and frozen at −80F. Nasha died in the summer of 2025 due to a gastrointestinal torsion.

### METHOD DETAILS

#### Data generation

##### DNA and RNA extractions

High-molecular-weight DNA was isolated either from whole blood using the UHMW Blood extraction protocol (Circulomics) or from testes tissue utilizing a HMW Phenol:Chloroform protocol.

RNA was isolated from blood using the Tempus™ Blood RNA Tube and Tempus™ Spin RNA Isolation Kit protocol (ThermoFisher) or from tissues utilizing a TRIzol Reagent (Invitrogen) and Qiagen RNAeasy protocol (Qiagen).

##### Pacific Biosciences HiFi long read sequencing

Pacific Bioscience (PacBio) HiFi libraries were generated for the mGirTip1. DNA was sheared using a Diagenode Megaruptor 3 to 20 kb mode size. At all steps, DNA quantity was checked on a Qubit Fluorometer with a dsDNA BR Assay kit (Thermo Fisher), and sizes were examined on a Fragment Analyzer (Agilent Technologies). SMRTbell libraries were then prepared for sequencing according to the protocol ‘Procedure-checklist-Preparing-whole-genome-and-metagenome-libraries-using-SMRTbell-prep-kit-3.0’. The libraries were then size-selected on the Blue Pippin using the BLF-7510 cassette under the 0.75% DF Marker S1 high-pass 15-20 kb protocol (Sage Science) to remove fragments below 15 kb in size. The selected library fractions were bound with the Revio Polymerase Kit and sequenced on a Revio instrument (PacBio) with 30 h movie time. 228 Gb of HiFi data was generated.

PacBio Iso-Seq libraries enable direct sequencing and analysis of full-length transcript isoforms. A PacBio Iso-Seq library was created using the ‘Preparing Iso-Seq® v2 libraries using SMRTbell® prep kit 3.0’ from the testes sample from the mGirTip1. The library was bound with the Sequel II Polymerase Kit and run on one Sequel II SMRT Cell in a 24-hour movie.

The PacBio Kinnex library preparation kits are based on the MAS-Seq method, which concatenates smaller amplicons into larger fragment libraries (Pacific Biosciences). Full-length RNA Kinnex libraries were made from RNA extracted from both the blood sample and the testes following the protocol ‘Preparing-Kinnex-libraries-using-the-Kinnex-full-length-RNA-kit.’ Kinnex libraries are pooled and bound with the Revio Polymerase Kit and run on a total of 7 Revio SMRT cells in 24-hour movies.

##### Oxford Nanopore long read sequencing

For the mGirTip1, a template for long read sequencing on the PromethION instrument was prepared using both the Ligation Sequencing Kit LSK-114 and the Ultra-Long DNA Sequencing Kit ULK-114 (Oxford Nanopore Technologies, Oxford, UK). For the LSK-114 kit, modifications to the DNA handling and cleanup procedure were utilized (https://community.nanoporetech.com/posts/rocky-mountain-adventures). 10 μg of DNA was extracted from blood and testes, cleaned and concentrated by addition of 1 volume of AMPureXP bead solution. After collecting the beads by magnet, the beads were washed 2 times with 0.5 mL 80% ethanol and dried for 1 min. The beads were suspended in a 51 μL elution buffer (EB) and DNA eluted at 37°C for 15 min. The sheared, eluted DNA was then processed by ligation to the ligation adapter (LA) as recommended by the manufacturer’s instructions, apart from an increase in the room temperature incubation of the ligation reaction to 60 minutes. Libraries were sequenced in 21 flow cells on a PromethION instrument using R10.4.1 flow cells. Two Ultra-Long DNA Sequencing Kit V14 (ULK-114) libraries were prepared, one from blood and one from testes tissue, following manufacturer protocols (Oxford Nanopore Technologies, Oxford, UK) and run on PromethION v10.4.1 flow cells.

##### Illumina short read sequencing

DNA from the mGirTip1 was then processed into a short read library using the TruSeq DNA PCR-Free Kit as recommended by the manufacturer (Illumina Inc., San Diego, CA) and sequenced on an Illumina NextSeq2000 platform using 2 × 151 base paired end reads. 147 Gb of total data was obtained.

An Illumina Stranded mRNA library was generated from testes tissues for the mGirTip1. Illumina Stranded mRNA libraries utilize a rapid, cost-effective workflow for accurate, unbiased detection of the protein-coding transcriptome with precise measurement of strand information (Illumina, San Diego, CA). The Illumina Stranded mRNA library was sequenced on the Illumina NextSeq2000 instrument (USDA MARC) with 2 x 101 base paired end reads.

To gather information about genome spatial organization for use in scaffolding, we generated a Dovetail Omni-C Hi-C library by cross-linking approximately 10 mg of testes tissue of mGirTip1. The Dovetail Omni-C Hi-C library was following the Dovetail Omni-C Protocol for Tissues v2.0 (Dovetail Genomics, Scotts Valley, CA). Omni-C libraries deliver uniform, shotgun-like sequence coverage that is utilized for enabling genotyping and haplotype phasing, alongside long-range information characteristic of Hi-C data. The library was sequenced on an Illumina NextSeq2000 sequencer with paired end 2 × 151 reads.

Whole-genome sequences of mGirTip3 and mGirTip2, the parents of mGirTip1, were generated using short-read Illumina sequencing. DNA for mGirTip3 came from post-mortem skin tissue and for mGirTip2 from tail hair root cells collected at the Cincinnati Zoo and Botanical Garden. Libraries were prepared at the Pennsylvania State University Huck Institute’s Genomic Core and sequenced at the Penn State Hershey Genomics facility on a NovaSeq using 150 bp paired-end reads. Demultiplexing was performed with Bcl2fastq, which masked short adapter-overlapping bases. Read quality was assessed with FastQC v0.11.8, showing high base quality across reads, and Fastp v0.20.0 confirmed that adapter trimming was unnecessary. Reads were aligned to the chromosome-level Masai giraffe genome assembly ASM165123v1 (Hi-C improved) using BWA-MEM (v0.7.17-r1188) with the -M flag. Alignments were filtered, sorted, and processed with Samtools, duplicates were removed with Picard MarkDuplicates, and final coordinate-sorted BAM files were evaluated for alignment and coverage statistics using Samtools stats and BEDTools.

##### T2T-mGirTip1v0.1 assembly

We assembled the giraffe genome with Verkko in trio mode using a combination of 90x PacBio HiFi and 45x Oxford Nanopore Technologies (ONT) duplex data (length > 20 kb, Q > 20) passed to Verkko as long-accurate reads (--hifi), 370x ONT simplex data (45x reads greater than 100 kb) passed to Verkko as ultra-long reads (--nano), and Illumina data (38x maternal, 72x paternal) from the parents for phasing. The assembly was generated with Verkko v2.2 using the parameter --unitig-abundance 4 due to the high depth of long-accurate reads used. Verkko was then rerun using the trio output and 200 million Hi-C read-pairs to assist in scaffolding across unresolved tangles in the graph.

#### Verkko-Fillet Functions

##### Verkko object

The Verkko object is a specialized Python class designed to consolidate extensive information into a single entity. This object can be easily saved and loaded from disk while maintaining a trackable history. It contains 11 key attributes, most of which are Pandas DataFrames^58^, making them easy to parse and use independently.

● .verkkoDir: Directory containing the verkko results.
● .verkko_fillet_dir: Output directory for verkko-fillet results.
● .paths: Path file extracted from assembly.paths.tsv.
● .version: Genome version.
● .species: Genome species.
● .stats: Assembly statistics (e.g., contig and chromosome information, completeness).
● .gaps: Gap information extracted from the .path file.
● .gaf: Graph alignment results (GAF format).
● .qv: Quality value (QV) calculation results using Meryl and Meruqury.
● .history: Log of all modifications made during the gap-filling process.
● .scfmap: Assembly scaffold mapping file (assembly.scfmap).

To generate the Verkko Fillet object from the Verkko result directory, the directory itself is the only required input.

`obj = vf.pp.read_Verkko(<VERKKODIR>)`

##### QV score

To estimate the base-level accuracy of the assembly, we calculated QV scores using a k-mer–based approach implemented with Meryl (v1.4.1). First, 31-mers were generated independently from Illumina and HiFi read sets. k-mers present more than once were extracted from both datasets, and a hybrid k-mer database was then constructed by combining the filtered Illumina and HiFi k-mers. This hybrid k-mer set was compared against the k-mers from assembly to identify k-mers absent from the read data, which were considered potential sequencing or assembly errors. QV scores were computed using Merqury (v1.3)^59^, which can also be performed via the Verkko-Fillet function.

`vf.tl.calQV(obj)`

##### Contig stats

To retrieve T2T statistics, including the number of scaffolds, contigs, telomeres, and gaps, the final assembly.fasta file from the Verkko directory is used.

`vf.tl.getT2T(obj)`

This function uses the `getT2T.sh` script from the marbl training github (https://github.com/marbl/training/blob/main/part2-assemble/docker/marbl_utils/asm_evaluation/getT2T.sh. Specifically, it employs seqtk gap to identify gaps and seqtk telo to detect telomeres at both ends of contigs, with CCCTAA as the telomeric sequence. In some cases, seqtk telo fails to detect telomeres that are not sufficiently close to contig ends. To improve detection, the function first runs seqtk telo -d 50000. It then runs seqtk telo -d 50000 again after trimming 2,000 bp from both ends using trimfq -b 2000 -e 2000. Finally, it runs seqtk telo -d 5000 after an additional 4,000 bp trim using trimfq -b 4000 -e 4000. The detected telomeric regions are merged using bedtools, and the output files are saved in the chromosome_assignment folder within the main Verkko-Fillet directory. These files identify T2T contigs and provide BED annotations for telomere and gap regions.

##### N50 and L50

The N50, which measures the contiguity of an assembly, is calculated using the contig statistics from obj.stats. To determine the N50, all contigs (or scaffolds) in the assembly are arranged in descending order by length. The N50 value is the length of the shortest contig for which the sum of all contigs of equal or greater length covers at least 50% of the total assembly length.

`vf.pl.n50Plot(obj)`

The function reports the N50 and L50 value and generates a bar plot representing sequence lengths. The N50 value is displayed and saved along with the plot.

##### Chromosome assignment

If a reference genome for the species exists, chromosome assignment for the assembly can be performed. This is achieved by aligning the assembly to the reference using Mashmap^60^.

`vf.tl.chrAssign(obj = obj, ref = ref)`

This function runs Mashmap using the command: mashmap -r <reference> -q <assembly> --pi 95 -s 10000 -t 8. The results are then filtered to retain segments with a block length greater than 1 Mbp and an identity of 99% for chromosome assignment.

`vf.pp.readChr(obj)`

The Mashmap alignment results can be read into obj.stats using the readChr function. The obj.stats table includes chromosome assignments, T2T contig statistics, and completeness data from the getT2T function.

##### Graph Alignment

To identify ONT reads that support path traversal across gaps for gap filling, we used GraphAligner^46^ to align ONT reads to the assembly graph.

`vf.tl.graphIdx(obj)`

The graph is indexed before running GraphAligner with --diploid-heuristic 21 31 and --diploid-heuristic-cache, which are heuristic parameters designed to better handle alignments in diploid genomes. Additional parameters related to seeding, extension, and indexing are also used to make this consistent with the Verkko assembler: --bandwidth 15, --seeds-mxm-length 30, --mem-index-no-wavelet-tree, and --seeds-mem-count 10000.

`vf.tl.graphAlign(obj, ontReadList = “ont.list”)`

Using the index generated above, ONT reads are aligned to the graph using GraphAligner. In this step, the diploid-related parameters --diploid-heuristic 21 31 are used again. Additionally, the following parameters are applied: --multimap-score-fraction 0.99, --precise-clipping 0.85, --min-alignment-score 5000, --hpc-collapse-reads, --clip-ambiguous-ends 100, --overlap-incompatible-cutoff 0.15, --max-trace-count 5, and --mem-index-no-wavelet-tree. This process generates a GAF (Graph Alignment Format) file for each ONT read with its corresponding paths.

`vf.pp.readGaf(obj)`

This function reads and parses the GAF file, storing it in the obj.gaf attribute for use in downstream analysis.

#### Align HPC node sequences to the assembly for IGV compatibility

To locate each node within the assembly—which can be useful during gap filling—we provide the script `_node_to_assembly.sh`. This script requires a path file, the HPC node sequences, the assembly, and the name of the target contig.

First, the list of nodes is extracted from the target contig’s path, and the corresponding HPC sequences are retrieved from the HPC node sequence set. Next, the sequence of the target contig is extracted from the assembly, and an index is generated with minimap2 (v2.30)^61^ using the -H option, which enables homopolymer-compressed k-mers. The HPC node sequences are then aligned to this index with minimap2, and the resulting BAM file is converted to BED format using bedtools(v2.31.1)^62^ bamtobed.

#### Gap filling

The path file, assembly.paths.tsv, generated by the Verkko assembler, contains three columns: contig name, path, and assignment. The gap information can be retrieved from the path column in this file. When the Verkko-Fillet object is created using the `read_Verkko()` function, the path file is stored in obj.path.

`vf.pp.findGaps(obj)`

The gap information includes flanking nodes and haplotype details, which helps prevent confusion between different gaps on the same chromosome and gaps with the same flanking nodes but on different haplotypes. This function extracts that information and stores it in obj.gaps.

`vf.pp.searchNodes(obj,[node list])`

To count the number of ONT reads that support walks spanning the nodes flanking a gap, this function creates a new attribute, obj.paths_freq, by parsing obj.gaf. By default, it filters the ONT reads to retain only the alignment with highest MAPQ per read, preventing duplicate counting of the same read. The function then reports the number of supporting reads for each walk (path), considering both directions.

`vf.pp.highlight_nodes(obj, node = <node>)`

In some cases, one of the nodes in a bubble may already be used in the other haplotype, and the coverage for both nodes may be sufficient to consider them as part of a haplotype. In such cases, we can assign the ambiguous node to the other haplotype, even if there is no ONT support showing a direct path through the ambiguous node with flanking haplotype-specific nodes.

`vf.pp.calNodeDepth(obj)`

To estimate the number of loops in repeat regions and exclude nodes from complex regions, it is useful to calculate node coverage. This can be done by analyzing obj.gaf, where the number of aligned reads per node is counted. The raw read counts are then normalized by the node length to obtain coverage values. This coverage information is merged into obj.node, allowing both haplotype assignment and coverage metrics to be assessed together for better-informed filtering.

`vf.pp.searchSplit(obj,node_list_input)`

If an edge between nodes is missing in the middle of a contig, it is possible to identify supporting ONT reads that align to both sides of the contig. To do this, the function filters for ONT reads that align with more than 5,000 bp at the ends of each contig (within 5,000 bp from the boundary) and have a mapping quality (MAPQ) greater than 0.

Verkko requires at least three supporting reads to create an edge. Therefore, if only two reads meet these criteria, Verkko may fail to generate the corresponding edge. This function helps recover such cases by identifying the supporting ONT reads, which are then incorporated into the final path file to improve connectivity.

`vf.pp.fillGaps(obj=obj, gapId= <gapid>>, final_path=<new path>).`

This function verifies the nodes and orientations of boundary elements to prevent incorrect gap filling. If discrepancies in node identity or orientation are detected at the boundaries, the function issues a warning.

Check the status and history of gap filling using `vf.pp.checkGapFilling(obj)`. There is also a function to delete a specific gap, `vf.pp.deleteGap(obj, gapId)`, to prevent incorrect fixes.

#### Detect internal telomere and trimming

Sometimes, artificial sequences extend beyond the true telomere, causing a contig to be incorrectly labeled as not T2T. This issue can be resolved by trimming these artifacts, provided they are confirmed to be non-biological.

`vf.tl.detect_internal_telomere(obj)`

This function runs the script originally used in the VGP assembly project ( https://github.com/VGP/vgp-assembly)^6^. Briefly, the script takes an assembly FASTA file as input and applies a distance threshold to define terminal and internal regions at both ends of each contig.

`result_merged, tel = vf.pp.find_intra_telo(obj, file=file, loc_from_end=15000)`

From the BED file generated by `vf.tl.detect_internal_telomere(obj)`, adjacent telomeric regions are merged. For each merged region, the highest proportion of telomeric sequence is retained. By default, only merged regions with a telomeric proportion greater than 0.5 are kept. Additional filters are applied: the total length of the merged region must be greater than 0 bp, and the region must start at least 15,000 bp from the contig boundary. This function returns two DataFrame, `<result_merged>` with internal and non-internal telomeres for both ends for each contig with other information such as the length of contig, or chromosome assignment result. `<result_merged>` is going to be used for plot heatmap of proportion of the telomere for each contig for both sides of contig using `vf.pl.percTel()` function.

`vf.pp.find_reads_intra_telo(tel, <line number>)`

Another DataFrame, tel, is generated by `vf.pp.find_intra_telo(obj)` and is used to extract reads covering the region preceding the internal telomere. This helps validate the structure and assess whether the region is a true biological signal or an assembly artifact. In some cases, reads in this region may originate from only one platform—such as ONT—even when the assembly was built using both HiFi and ONT data. The function locates the relevant contig segment and extracts the corresponding reads based on their relative position in HPC space. To visualize the region, the `vf.pl.readOnNode(tel, <line number>, df)` function can be used.

`vf.tl.runTrimming(obj, <dict>)`

If a dictionary in BED file format is provided, with keys for contig, from, and to, this function trims the assembly (by default, assembly.fasta) to the specified region and saves the trimmed version with the suffix “_trimmed.fasta”.

#### Finding broken contigs and connecting them

Not only gaps, but also broken contigs need to be correctly connected by identifying their counterpart contigs. There are two scenarios that require contig connection: first, when a contig is broken into two pieces, but both fragments are long enough to be assigned a haplotype and chromosome; second, when one of the fragments is too short to be assigned a chromosome, necessitating its connection with the longer fragment.

##### Scenario 1: Both contigs can be assigned to the same haplotype and chromosome

`vf.pp.detectBrokenContigs(obj)`

All T2T contig statistics, including haplotype and chromosome assignments for the main contigs, are stored in obj.stats. The function uses this information to identify pairs of contigs that belong to the same haplotype and chromosome. It then determines which chromosomes should be connected.

##### Scenario 2: One of the contigs is not able to be assigned a chromosome

In this scenario, the sequences of nodes are aligned pairwise to identify the counterpart nodes on the other haplotype, allowing us to determine which contigs should be connected.

`vf.tl.gfaToFasta()`

We extracted the HPC (homopolymer-compressed) sequences of each node from the GFA^41^ file into a FASTA format. By default, the file assembly.homopolymer-compressed.gfa is used, and the resulting FASTA file is saved with the same prefix, but with a .fasta extension.

`vf.tl.mapBetweenNodes()`

The FASTA file containing node sequences is aligned pairwise using Mashmap^60^, with self-alignments skipped by default.

`vf.pl.nodeMashmapBlockSize(node = <node>)`

The Mashmap output is parsed and used to sort all nodes aligned to a specific `<node>` (used as the query), based on the total aligned block size. The block sizes are grouped and summed by reference nodes to rank the best-matching counterparts.

#### Connect the two contigs to each other

`vf.pp.connectContigs(obj, contig = <sor>, contig_to = <target>, gap = <gapid>, at = >left of right>, flip = <false>)`

After identifying the two contigs that should be connected, either contig can serve as the source or the target. At this stage, the orientation and order of the contigs are critical and must be carefully verified—preferably by inspecting the graph in Bandage^44^—before proceeding with the connection. This function does not modify obj.paths directly. Instead, it adds a new gap record to obj.gaps, containing the source and target contig names, their orientations, and connection order. These entries can later be processed by the `vf.pp.updateConnect(obj)` and `vf.pp.writeFixedPaths(obj)` functions, as described below.

#### Clean up the paths

After gap filling with the Verkko-Fillet module, the original paths must be updated to incorporate the newly resolved sequences. This process involves not only replacing gaps with the filled segments but also updating connections between nodes to reflect the improved contiguity.

To avoid misassignment of reads to irrelevant or unused nodes during the Verkko consensus step, it is recommended to filter out certain paths. Specifically, short paths composed only of disconnected nodes, nodes used exclusively in gap filling, or nodes that were never part of the original assembly should be excluded.

`obj = vf.pp.keepContig(obj, contig_lst, path_lst)`

By default, this function utilizes the list of contigs stored in obj.stats. However, if key contigs are missing—such as mitochondrial sequences or other small scaffolds—they should be added manually at this stage. The obj.paths DataFrame is then updated with a new column, ’rm’, which logs filtering decisions. Contigs designated for retention are labeled “keep_contig”.

`obj = vf.pp.checkDisconnectNode(obj, min_hpc_len=100_000)`

When two contigs are connected using the `vf.pp.connectContigs()` function, the changes are recorded in obj.gaps, but not yet reflected in obj.paths. To finalize these connections, this function marks the source and target contigs within obj.paths. This annotation allows `vf.pp.writeFixedPaths(obj)` to correctly merge them by filtering out the target contig and connecting it with the source, preparing the structure for consensus generation.

`obj, cludlst = vf.pp.keepNodesInUnresolvedGaps(obj)`

Some nodes may not have been utilized in the final assembly but remain connected to paths without gaps. This typically occurs when the assembly graph contains many nodes, but tools like Rukki select only a subset for the final paths. These unused nodes can still attract reads, potentially diverting them from the correct locations.

To address this, a directed graph is constructed from obj.edges using NetworkX^63^. The graph is traversed to identify all nodes upstream and downstream of each gap using nx.ancestors() and nx.descendants(). For gaps between newly connected contigs, edges linking the source and target are added. Nodes found within unresolved gaps are flagged with “keep_Nodes_in_unresolved_gaps” in the final path annotation.

`path = vf.pp.writeFixedPaths(obj)`

Once all filtering steps are complete, the original paths are replaced with the refined, gap-filled versions. Connections between source and target contigs are finalized, and redundant or trivial paths—especially those with nodes already used in gap filling—are removed. The finalized path file and corresponding GAF file are then written to disk, ready for the next step in the Verkko consensus workflow.

### Run Verkko consensus using the final path with filling the gaps

Verkko automatically determines which steps to run by checking the presence and timestamps of relevant files. Therefore, it is important to ensure that the working directory is properly set up with all necessary files before running the consensus step. Additionally, if new edges are added, several files must be updated to include the corresponding supporting ONT reads and walks to ensure accurate consensus generation.

`vf.pp.mkCNSdir(obj, <new_folder_path>, missingEdge=<True or False>)`

This function generates a new folder, enabling Verkko to resume from the consensus step while incorporating any new nodes and edges introduced by the user during this stage.

Verkko --slurm -d <new_folder_path> [ --hifi <hifi reads> --nano <ont reads> ] --screen rDNA <rDNA.fasta> --screen mitochondria <mitochondrial.fasta> --paths <final path file> --assembly <original verkko directory> [--snakeopts "--dry-run"]

Since version 2.3, Verkko automatically inherits the read sets from the existing assembly, so this command can be simplified by removing --hifi and --nano.

To generate cleaner results and remove redundant sequences from rDNA, we used rDNA sequences and giraffe mitochondrial sequences (MT605038.1.fasta) to screen mitochondrial nodes. Before running Verkko, it is advisable to use the `--snakeopts “--dry-run”` parameter with the Verkko consensus command. This will allow you to verify that Verkko is only submitting jobs for `buildPackages, cnspath, combineConsensus, and layoutContigs.`

### Tuning and polishing

#### Naming the unplaced nodes

The unplaced sequences were assigned chromosome information by leveraging the edges in the graph. Using the edge and node data from the GFA^41^, we clustered connected nodes together and assigned chromosome labels based on nodes that could be reliably mapped to a chromosome. Prior to this step, we excluded nodes that were used multiple times across different chromosomes to prevent ambiguity.

`duplicate_nodes, node_database = vf.pp.find_multi_used_node(obj)`

This function lists all nodes from each path that has been assigned a chromosome in obj.stats and groups them by chromosome to create a clean list of nodes for each chromosome. It then identifies nodes that are used across two different chromosomes. For example, if node utig4-0 is used in both path_number_1 assigned to chr1 and path_number_2 assigned to chr2, the function will return utig4-0 as a “multiple-used” node to exclude it from naming with `vf.pp.naming_contigs()`. For the sex chromosomes, chrY is treated as chrX to prevent mislabeling of nodes in the PAR region as multiple-used. The function returns duplicate_nodes and node_database, which is a table containing chromosome information and the list of nodes from each path.

`chrMap = vf.pp.naming_contigs(obj, node_database, duplicate_nodes)`

Using the outputs from the previous function, this function reads the GFA^41^ to obtain edge information and constructs an undirected network using the Python NetworkX package^63^. It then clusters the nodes based on their connections while excluding nodes that are marked as “multiple-used.” This process results in a clean cluster of fully connected nodes. Chromosome information is assigned to all nodes within each cluster if the chromosome information from any node in the cluster is unique. This allows us to determine the chromosome assignment for unplaced nodes. The output, chrMap, is a table containing the contig names from the assembly, along with their assigned chromosome and haplotype names.

`vf.tl.renameContig(obj, chrMap)`

The renaming of all contigs in the assembly is performed using this function. The output FASTA file is saved with the suffix `_sorted.fasta` appended to the input assembly file name.

#### Flipping and sorting assembly

The assembly may have the reverse orientation compared to the existing reference or the reference for orthologous species. Before finalizing the assembly, it is recommended to align its orientation with that of the widely used reference and ensure it is properly sorted.

`vf.tl.flipContig(<list of contig to be flipped>)`

The assembly.fasta file in the Verkko-Fillet directory is used, and the output is saved with the suffix _flipped.fasta in the same directory.

`vf.tl.sortContig(sort_by = “hap”)`

There are two options for the sort_by parameter: one based on haplotype (sort_by = “hap”) and the other based on chromosome number (sort_by = “chr”). The default option is hap.

#### Polishing

Assembly errors were corrected using the variant calling method described by McCartney et al^14^. and Yoo et al^50^. To do this, we constructed a haploid reference that included haplotype contigs along with chrX, chrY, and chrM, resulting in T2T-mGirTip1v0.2-Mat and T2T-mGirTip1v0.2-Pat. Illumina and HiFi reads were aligned to both haploid and diploid assemblies, whereas ONT reads were aligned only to the diploid assembly.

Illumina reads were aligned with bwa^64^ and then deduplicated using samtools^65^ markdup. ONT and HiFi reads were aligned with winnowmap (v2.03)^66^, using the -ax map-ont and map-pb parameters, respectively. Primary alignments were retained by filtering with samtools view - F0x104. Finally, BAM files from HiFi and Illumina were merged for each reference with samtools merge, hereafter referred to as the hybrid.

Hybrid alignments on haploid assemblies were processed with DeepVariant (v1.6.1)^67^ using the HYBRID_PACBIO_ILLUMINA mode, with a MAPQ threshold of 1 for BAM files aligned to haploid assemblies and 0 for those aligned to the diploid assembly. Variant calling from ONT alignments was performed with ONT_R104, applying a MAPQ threshold of 0.

VCF files from hybrid alignments on T2T-mGirTip1v0.2-Mat, Pat, and diploid (Dip) assemblies, as well as ONT alignments on T2T-mGirTip1v0.2-Dip, were passed to the snv_candidates.sh script^50^. The script was run with the hybrid k-mer database and an estimated haploid peak of 68, derived from the hybrid k-mer multiplicity distribution, to generate the final VCF file. Polishing variants were then applied to produce version T2T-mGirTip1-v0.2.4, excluding variants on chrM, using bcftools^68^ consensus -H.

#### rDNA patching

Even after generating the consensus with corrected variants, a high density of error k-mers remained within one rDNA unit on the chr14 paternal haplotype. To address this, we removed that unit and joined the two flanking units at the start site of the 45S alignment between consecutive units. This was done by creating three separate FASTA files: each was broken at the start site of the 45S alignment of the high-density error k-mers and the start site of the following unit, after which the first and last files were connected. The resulting assembly is T2T-mGirTip1v1.0-Dip.

#### Mitochondral DNA circularization

Three mitochondrial contigs were identified in T2T-mGirTip1v0.2-Dip using the screen-assembly.pl script from the Verkko library. The longest contig was then circularized with the circularize_ctgs.py script, also from the Verkko library. This final circularized mitochondrial contig was included in the submitted genome.

### QUANTIFICATION AND STATISTICAL ANALYSIS

#### Assembly assessment

##### NucFlag

The nucleotide frequency plots and genome misassembly region file were generated using NucFlag (v0.3.3)^69^ with HiFi alignments on the T2T-mGirTip1-v0.3 genome, applying default parameters (Figure 5A).

##### BUSCO

For each assembly, we assessed completeness using BUSCO (v6.0.0)^51^ with the cetartiodactyla_odb10 lineage dataset. To compare BUSCO scores between T2T-mGirTip1-v0.1 and v1.0, chrX, chrY, and chrM were also included in each haplotype assembly.

##### Benchmark using read alignment

Read alignments on T2T-mGirTip1-v0.1 were performed using the same read types and corresponding parameters described in [Polishing in Method]. Alignment quality statistics were obtained with Picard (v3.3.0, http://broadinstitute.github.io/picard) CollectAlignmentSummaryMetrics for each BAM file using default parameters.

We also compared alignment statistics between ASM1759144v1, T2T-mGirTip1-v1.0, and the primary assembly. Illumina and ONT data were downloaded from the NCBI database (accession ID PRJNA627604)^52^ and aligned to the corresponding references using the same parameters as for the T2T-mGirTip1 alignment. Quality was then assessed with Picard CollectAlignmentSummaryMetrics.

##### Flagger

We used the read-mapping-based tool Flagger (v1.1.0)^4^ to detect mis-assemblies in dual or diploid genome assemblies. The Singularity image was downloaded from the Flagger GitHub repository, and the tool was executed with Singularity (v4.2.2) using ONT and HiFi alignments on T2T-mGirTip1-v0.1 and T2T-mGirTip1-v1.0. The workflow consisted of two steps: (1) converting BAM files to COV format, and (2) detecting abnormal coverage.

For the first step, we used the following command, where platform (pf) represents either HiFi or ONT:

`singularity exec flagger.sif \ bam2cov --bam ${pf}_to_dip.pri.bam \`

`--output ${pf}_coverage_file.cov.gz \`

`--runBiasDetection \`

`--annotationJson annotations_path.json \`

`--baselineAnnotation genome.size`

For the second step, we used the following command, with parameters adjusted according to pf. For ONT, we specified

`alpha = alpha_optimum_trunc_exp_gaussian_w_8000_n_50.ONT_R1041_Dorado_DEC_2024 .v1.1.0.tsv`

with winlen = 8000. For HiFi, we specified

`alpha = alpha_optimum_trunc_exp_gaussian_w_16000_n_50.HiFi_DC_1.2_DEC_2024.v1. 1.0.tsv`

with winlen = 16000. Both alpha TSV files were downloaded from the HMM-Flagger GitHub repository (https://github.com/mobinasri/flagger).

`singularity exec flagger.sif \ hmm_flagger \`

`--input ${pf}_coverage_file.cov.gz \`

`--outputDir ${pf}_hmm_flagger_outputs \`

`--alphaTsv ${alpha} \`

`--labelNames Err,Dup,Hap,Col \`

`--windowLen ${winlen}`

##### Hi-C alignment

Hi-C data from ASM1759114v1 were downloaded from the NCBI database (accession ID PRJNA627604)^52^. To assess mapping quality, Hi-C reads from ASM1759114v1 and mGirTip1 were aligned to both T2T-mGirTip1v1.0-Pri and ASM1759144v1 using bwa mem -5SP (v0.7.17)^64^. The resulting SAM files were converted into pairs format using pairtools (v1.1.0)^70^ with sorting and deduplication. A mapping quality threshold of 1 was applied for the haploid assembly (ASM1759114v1 and T2T-mGirTip1v1.0-Pri).

The commands used for this analysis are provided below:

`bwa mem -t $thread -SP $ref $hic1 $hic2 | \`

`pairtools parse -c $ref.size --min-mapq $mapq |\`

`pairtools sort |\`

`pairtools dedup \`

`--output $asm.nodups.pairs.gz \`

`--output-dups $asm.dups.pairs.gz \`

`--output-unmapped $asm.unmapped.pairs.gz \`

`--output-stats $asm.dedup.stats`

Contact matrices were generated from deduplicated pairs files using cooler (v0.10.4)^71^. For each assembly ($asm), we loaded pairs into a single-resolution .cool at bin size $bin using cooler cload pairs, then produced a multiresolution .mcool with cooler zoomify at 100 kb, 500 kb, 1 Mb, and 2 Mb. Zoomified matrices were balanced during processing.

`cooler cload pairs \`

`-c1 2 -p1 3 -c2 4 -p2 5 \`

`--assembly $asm \`

`$ref.size:$bin \`

`$asm.nodups.pairs.gz \`

`$asm.$bin.cool`

`cooler zoomify \`

`--nproc $thread \`

`--out $asm.$bin.mcool \`

`--resolutions 100000,500000,1000000,2000000 \`

`--balance \`

`$asm.$bin.cool`

### Genome annotation

#### Non-syntenic regions

We mapped T2T-mGirTip1-v1.0 (query) to ASM1759144v1 (reference) using Mashmap (v3.1.1)^60^ with a segment length of 50 kb and a minimum percent identity of 99%. Mashmap output was converted to BED, restricted to chromosome-consistent matches, and subtracted from the T2T-mGirTip1-v1.0 whole genome length BED to identify non-syntenic regions.

`mashmap -r $ref \`

`-q $query \`

`--pi 99 \`

`-s 50000 \`

`-o ASM1759144v1.T2T-mGirTip1-v0.1.syntenic.mashamp`

`awk ’{print $1,$3,$4,$6,0,$5}’ OFS=’\t’ \ ASM1759144v1.T2T-mGirTip1-v0.1.syntenic.mashamp \`

`­ ASM1759144v1.T2T-mGirTip1-v0.1.syntenic.mashamp.bed`

`awk ’{print $1,$0}’ OFS=’\t’ \`

`ASM1759144v1.T2T-mGirTip1-v0.1.syntenic.mashamp.bed |\ sed -e ’s/_.at//’ |\`

`awk ’$1==$7 {print $2,$4,$5}’ OFS=’\t’ \`

`> ASM1759144v1.T2T-mGirTip1-v0.1.syntenic.mashamp.chr_match.bed`

`bedtools subtract -a assembly_dim.fasta.bed -b \ ASM1759144v1.T2T-mGirTip1-v0.1.syntenic.mashamp.chr_match.bed \`

`> ASM1759144v1.T2T-mGirTip1-v0.1.non.syntenic.mashamp.chr_match.bed`

#### Repeat annotation

The repeats in T2T-mGirTip1v1.0-Dip were annotated using RepeatMasker (v4.2.1)^72^ with Dfam (v3.9) and Giraffa tippelskirchi as the reference lineage. The resulting annotation table was converted into BED format using the RM2Bed.py script(https://github.com/Dfam-consortium/RepeatMasker). To calculate the percentage and total length of repeats within non-syntenic regions, we intersected the repeats identified by RepeatMasker with the non-syntenic regions using bedtools^62^.

#### Gene annotation

Gene annotation on the T2T-mGirTip1v1.0 was performed with EviAnn software (v2.0.3)^56^. EviAnn software produces evidence-based genome annotation. It derives gene annotations from aligned and assembled transcriptome sequencing data and protein homology. Illumina RNA-seq and PacBio Kinnex read data from testes tissue of mGirTip1 and cardiac muscle tissue of mGirTip4 were used as transcriptomic evidence, in addition to the blood of mGirTip1 sequenced with PacBio Kinnex. A set of 576,821 mammalian proteins was used as protein homology evidence. Function annotations of protein coding genes were performed by aligning protein sequences to the UniProt-Swissprot database^57^ of functional proteins. Table 2 shows counts of protein coding and long non-coding RNA genes, transcripts and proteins for both primary and alternative haplotype assemblies. 98% of all annotated transcripts were assigned function with alignments to UniProt.

Both the genome and the annotation were evaluated for completeness of the gene content using BUSCO^51^ version 5.8.3 with “mammalia” orthologous protein database containing 9,226 amino-acid sequences. BUSCO scores for the genome and the annotated proteins are shown in Table S4. Primary and alternative haplotype assemblies show excellent completeness, missing only 0.9% and 3.1% of BUSCOs. Note that while annotation has fewer fragmented BUSCOs (0.4%) compared to the assemblies (0.8%), it has about 0.8 percentage points more missing BUSCOs, which implies that BUSCO software was able to identify homologous matches in the genome, but were not able to identify such matches in the protein sequences.

#### PAR regions

We aligned chrY_pat to chrX_mat using minimap2 with the asm20 preset to identify pseudoautosomal (PAR) regions on the sex chromosomes. The resulting PAF file was filtered to retain long, high-identity alignments (≥10 kb and ≥95% identity), representing PAR homologous segments. These intervals were converted to BED format, retaining only the chrY-derived coordinates, used to mask the corresponding regions on the primary assembly with bedtools maskfasta, producing a PAR-masked version of the reference genome.

To match the orientation of chromosome Y with other ruminant genomes, such as cattle and sheep, we inverted the chromosome so that it begins with the large PAR region followed by the centromere, ensuring that the short arm appears first for consistency.

