## Supplementary material for "Finishing a complete giraffe genome from telomere to telomere with Verkko-Fillet": Figure S

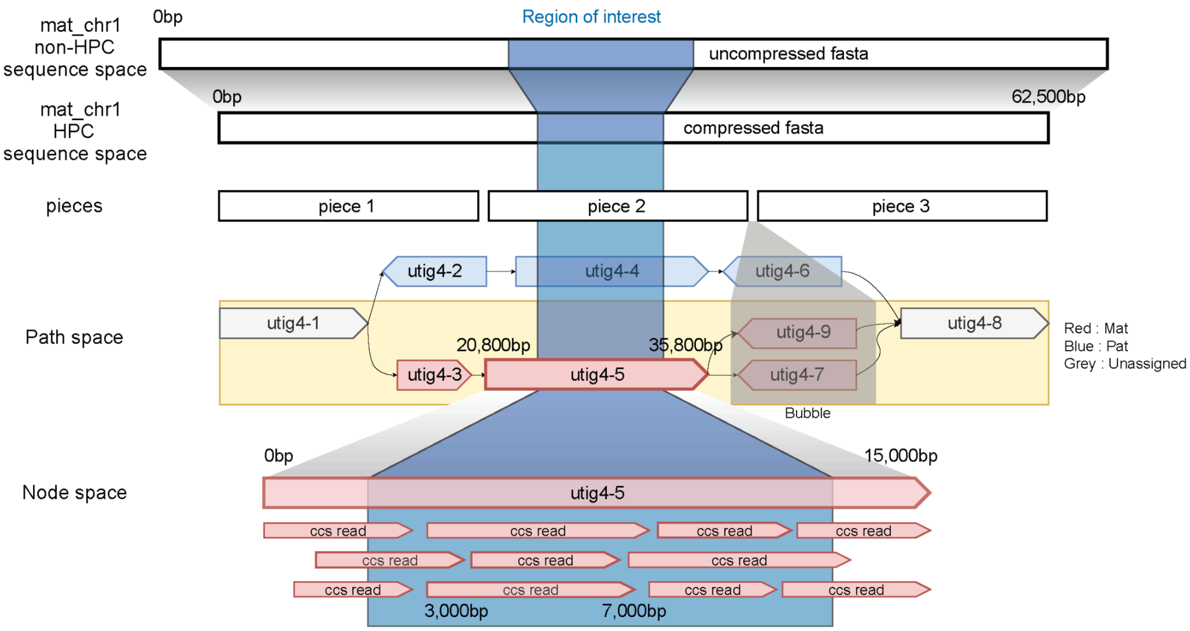


**Supplementary Figure 1. Verkko assembly structure.**

The Verkko assembly is organized into multiple layers. At the bottom is the node space, which records the reads used and later contributes to the consensus. Above this is the path space, where the order, connectivity, and orientation of nodes define how contigs are generated. For example, the maternal (red) haplotype is assembled from the path utig4-1(+), utig4-3(+), utig4-5(+), [gap], utig4-8(+). The next layer is the piece space, where paths are broken when the node walk becomes ambiguous and are represented as separate pieces, here between utig4-5 and utig4-8. Finally, all pieces are assembled into a homopolymer-compressed (HPC) sequence, concatenated with gaps, in which consecutive identical bases are collapsed into one. This compression is later reversed in the final non-HPC consensus assembly

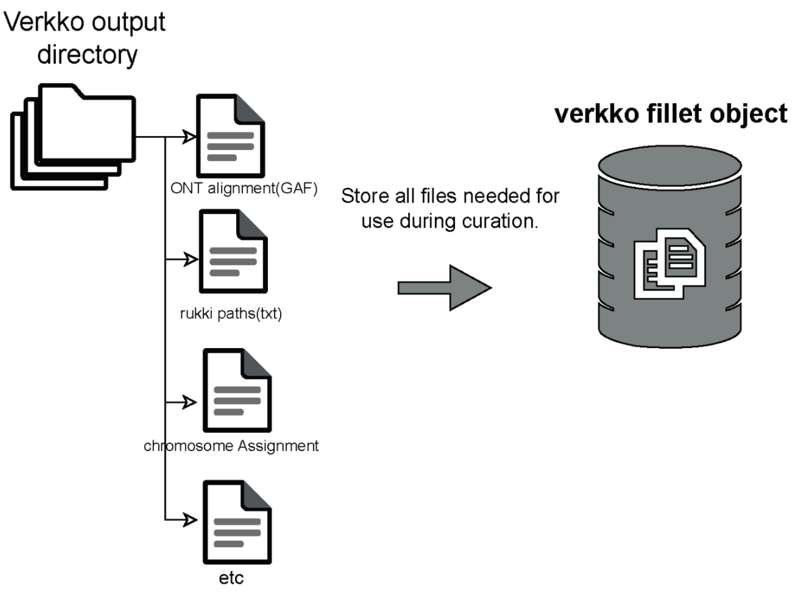


**vf-obj.gaps**

| gapId | contig | gap | fixdPath |
|---|---|---|---|
| gapid_0 | chr5_mat | [utig4-2639+, [N5000N:ambig_path], utig4-2652+] | utig4-2639+, utig4-2652+ |
| gapid_1 | chr5_mat | [utig4-70+, [N212780N:tangle], utig4-100+] | utig4-70+, utig4-99-, utig4-96-, utig4-97+, utig4-96-, utig4-98+, utig4-100+ |
| gapid_2 | chr5_mat | [utig4-100+, [N5000N:ambig_path], utig4-2421+] | utig4-100+, utig4-2421+, utig4-2421+, utig4-2421+ |
| gapid_3 | chr11_mat | [utig4-1130-, [N5000N:ambig_path], utig4-1082+] | utig4-1130-, utig4-1131+, utig4-333-, utig4-331+, utig4-334+, utig4-1087-, utig4-1083-, utig4-1082+ |
| gapid_4 | chr11_mat | [utig4-370+, [N100000N:scaffold], utig4-2178+] | utig4-370+, utig4-373+, utig4-2179-, utig4-2176-, utig4-2178+ |
| gapid_5 | chr10_mat | [utig4-1957+, [N5000N:ambig_path], utig4-2581-] | utig4-1957+, utig4-1961+, utig4-2583-, utig4-2581- |

**vf-obj.node**

| node | len | mat-kmer | pat-kmer | hap | ONT support |
|---|---|---|---|---|---|
| utig4-626 | 910848 | 29 | 36 | ambiguous | 10682 |
| utig4-629 | 3058 | 0 | 0 | ambiguous | 225 |
| utig4-630 | 8464 | 0 | 16 | ambiguous | 195 |
| utig4-2613 | 525203 | 6 | 28 | ambiguous | 6049 |

**vf-obj.paths**

| contig | path |
|---|---|
| chr10_mat | utig4-1795+,utig4-2059-,utig4-2060+, …, ,utig4-1672-,utig4-294-,utig4-290-,utig4-292+ |
| chr10_pat | utig4-2062-,utig4-2059-,utig4-2061+, … , ,utig4-1672-,utig4-293-,utig4-290-,utig4-291+ |

**Supplementary figure 2. The structure of the Verkko-Fillet object.**

The Verkko-Fillet object parses key outputs from the Verkko assembly directory, including paths, gaps, nodes, edges, and haplotype information. Gap filling is performed by integrating multiple data sources, such as ONT, HiFi, and Hi-C alignments. All attributes stored in the Verkko-Fillet object can be accessed in a dataframe format. In this figure, we show the gaps, nodes, and paths attributes, which represent key components of the Verkko-Fillet.

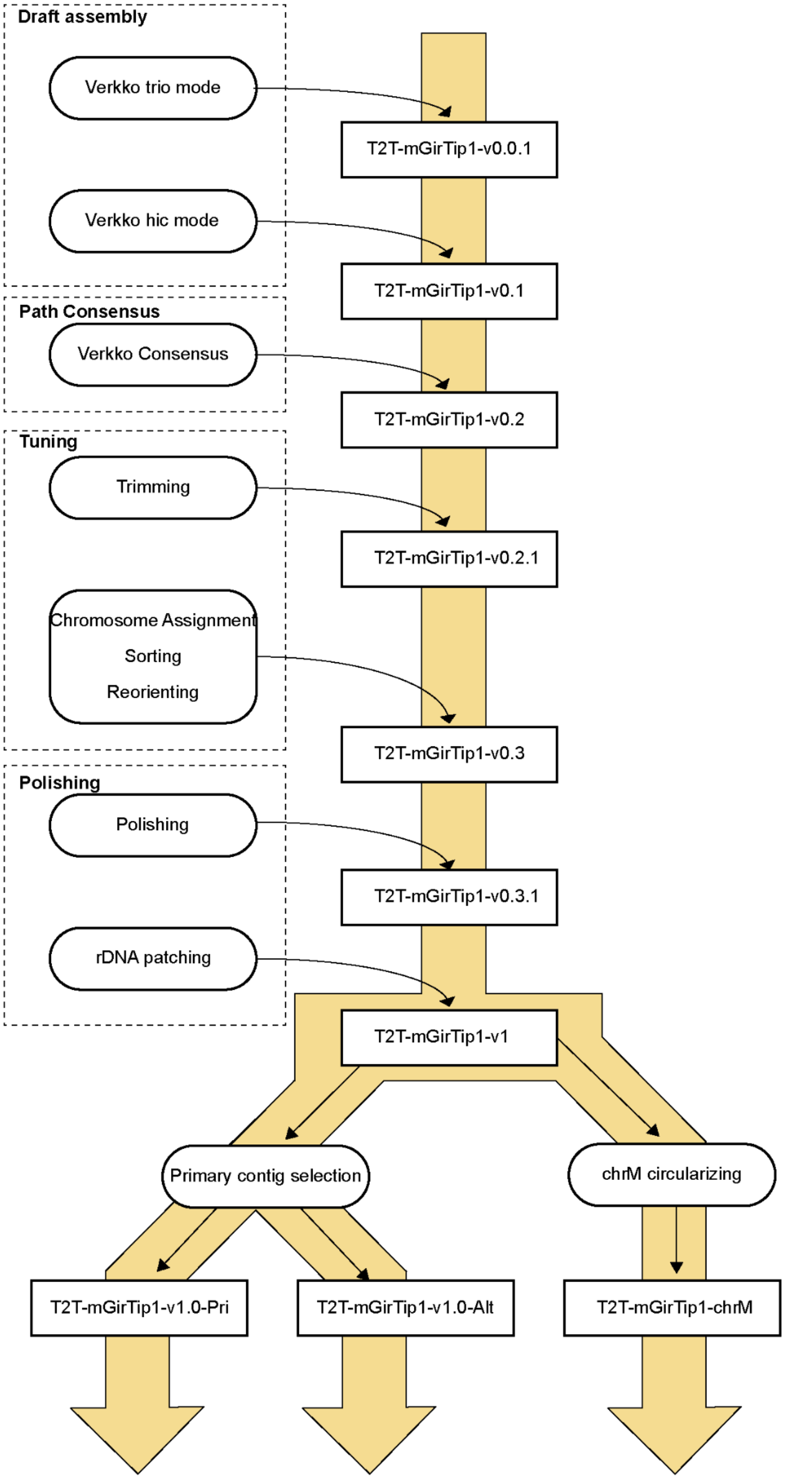


**Supplementary Figure 3. Versioning of T2T-mGirTip1 and associated workflow.**
The specific version shown here follows the workflow described in Figure 1. The final primary, alternative, and circularized mitochondrial DNA assemblies are provided as FASTA files, ready for genome submission.

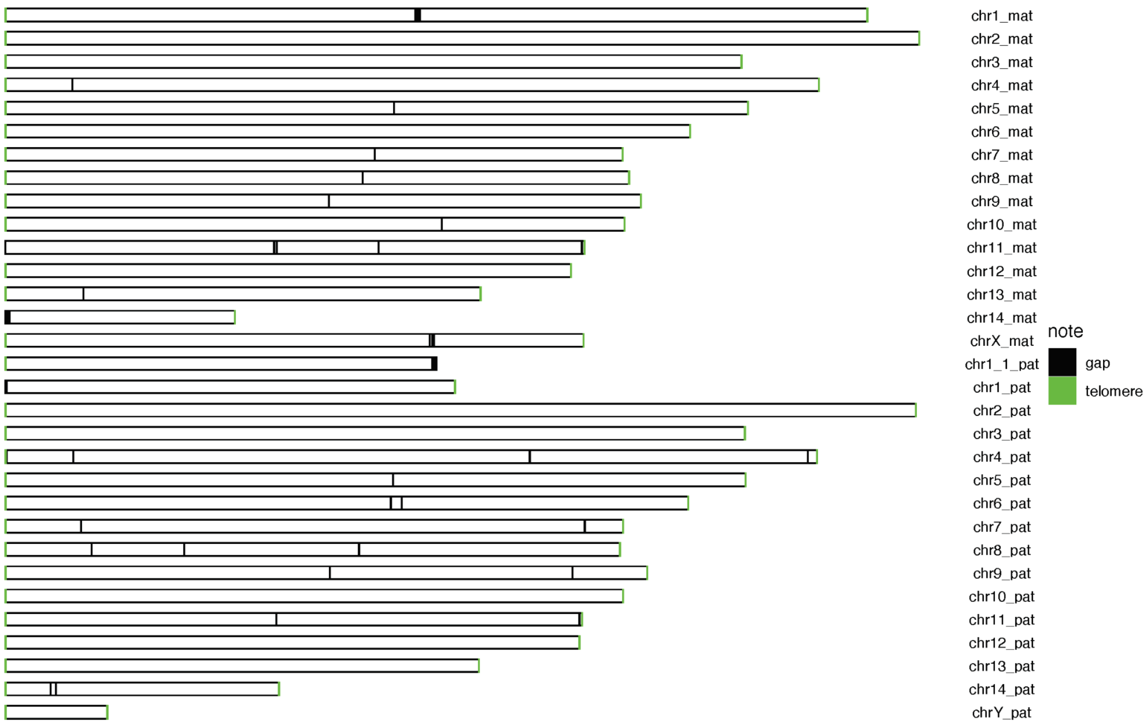


**Supplementary Figure 4. Ideogram of T2T-mGirTip1v0.1 with gap and telomere annotations.**

Black indicates gaps, and green marks telomere regions identified in T2T-mGirTip1v0.1. Chr. 1 paternal haplotype consists of two separate contigs (chr1_1_pat and chr1_pat), with only one telomere present at each contig end, and is nearly half the size of chr1 of the maternal haplotype.

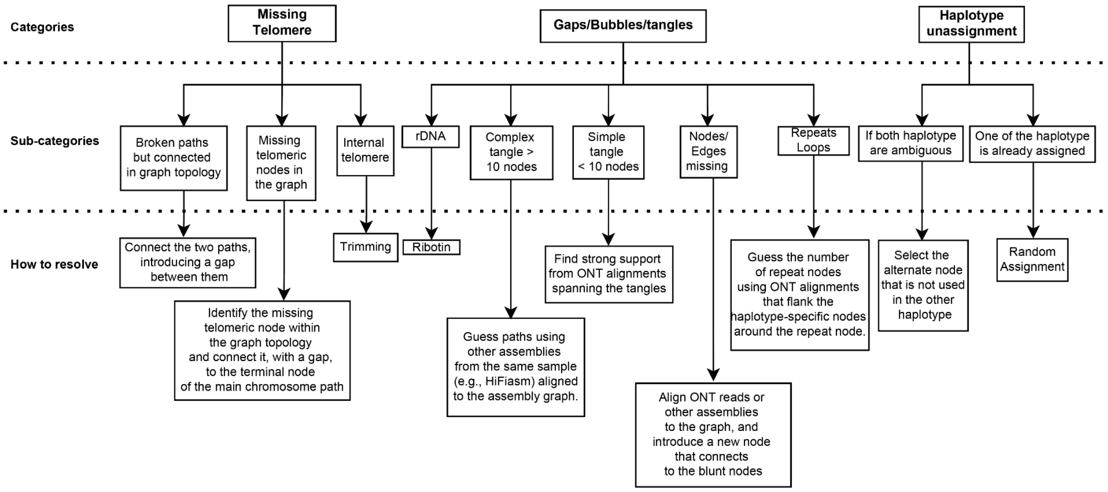


**Supplementary Figure 8. Decision-making tree for resolving tangles or correcting contigs.**

We defined three major categories—missing telomeres, tangles, and haplotype unassignment—each with their own subcategories. For every subcategory, we outline the corresponding strategy used to resolve the issue, providing a systematic framework for addressing assembly errors.

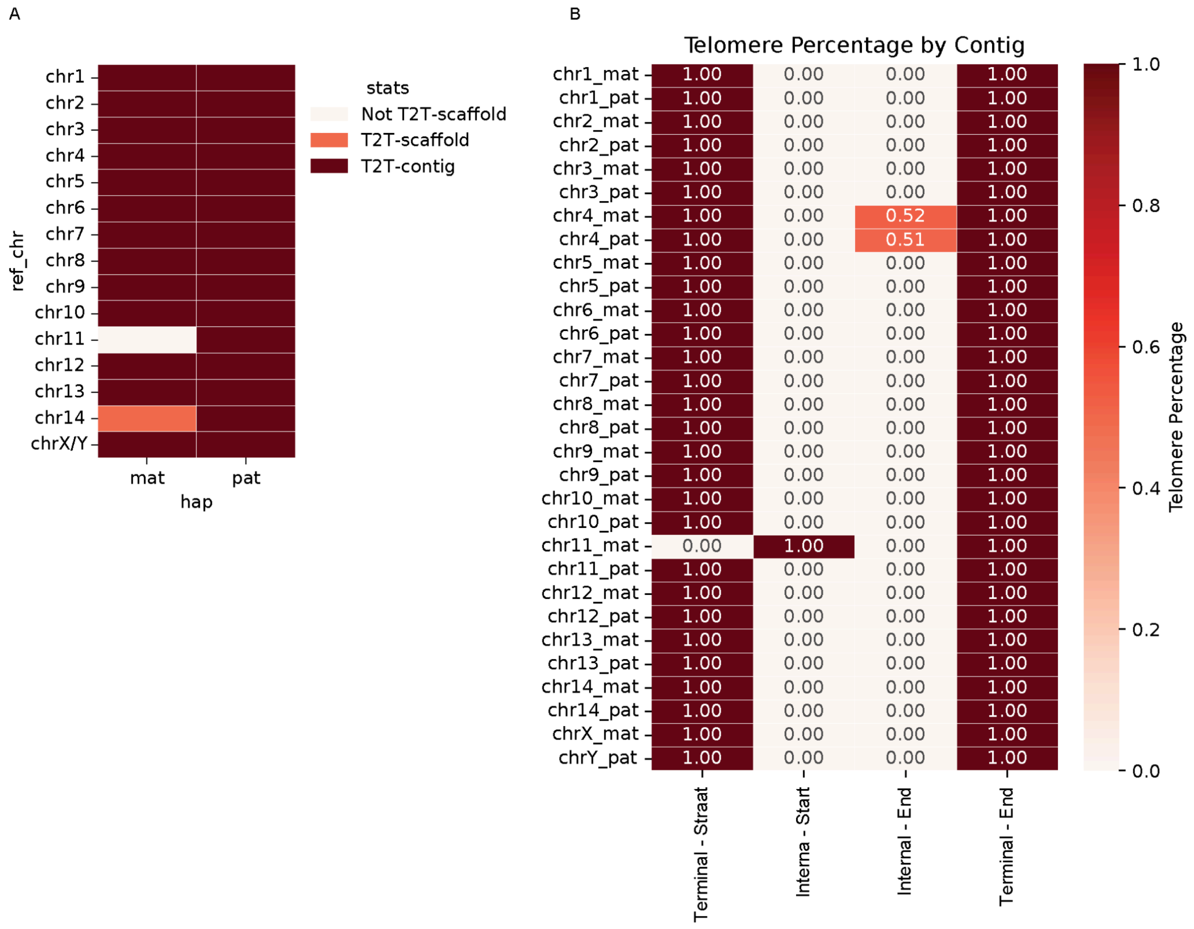


**Supplementary Figure 6. T2T statistics and telomere distribution in T2T-mGirTip1v0.1 (after gap filling).**

**(**A) After path correction, including gap filling and contig joining, one contig (chr11_mat) lacks a telomere at its end. Another contig (chr14_mat) is the only T2T scaffold, with telomeres at both ends but containing an internal gap. (B) Chr11_mat shows a high proportion of telomeric sequences internally.

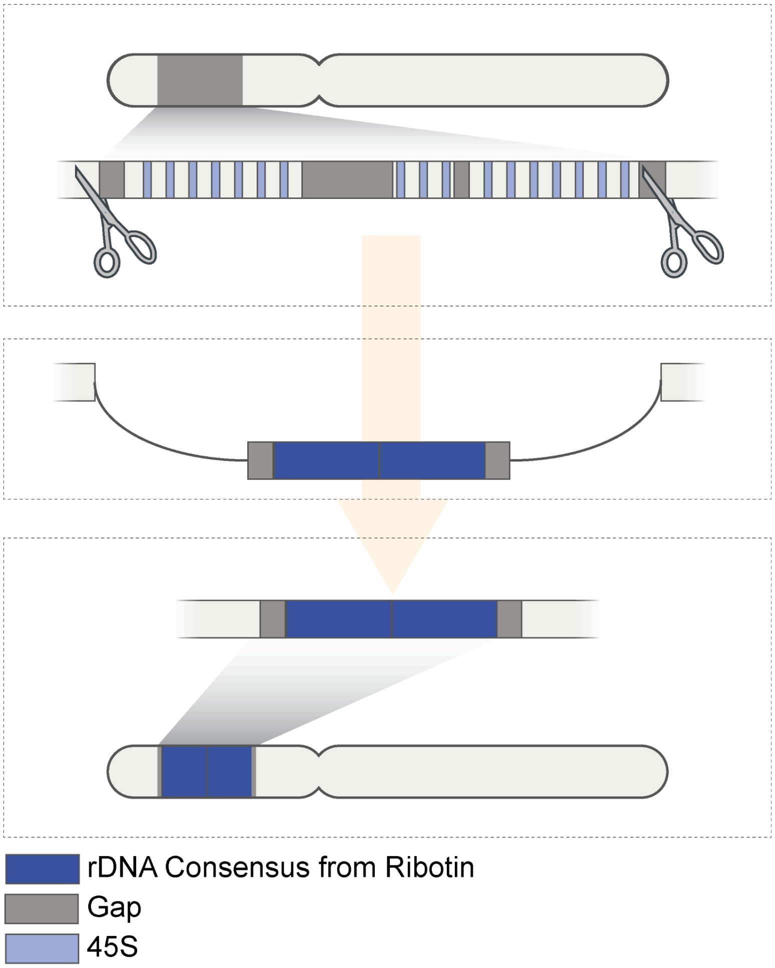


**Supplementary Figure 7. rDNA patching feature in Verkko-Fillet using rDNA consensus.** The workflow illustrates the process of patching rDNA regions. rDNA clusters often appear as mosaics of gaps and sequences, sometimes with multiple gaps within a single region. To address this, we provide a workflow that replaces the entire problematic rDNA cluster on a contig with a set of rDNA consensus sequences generated by Ribotin. In this case, the rDNA consensus sequences were derived from rDNA nodes within the graph. This approach ensures that intact rDNA units are included, which can be particularly valuable for genome alignment in downstream analyses. Because the exact copy number of the complete rDNA unit is not known, two gaps remain flanking the inserted consensus sequence—between the rDNA consensus and the original contig sequence.

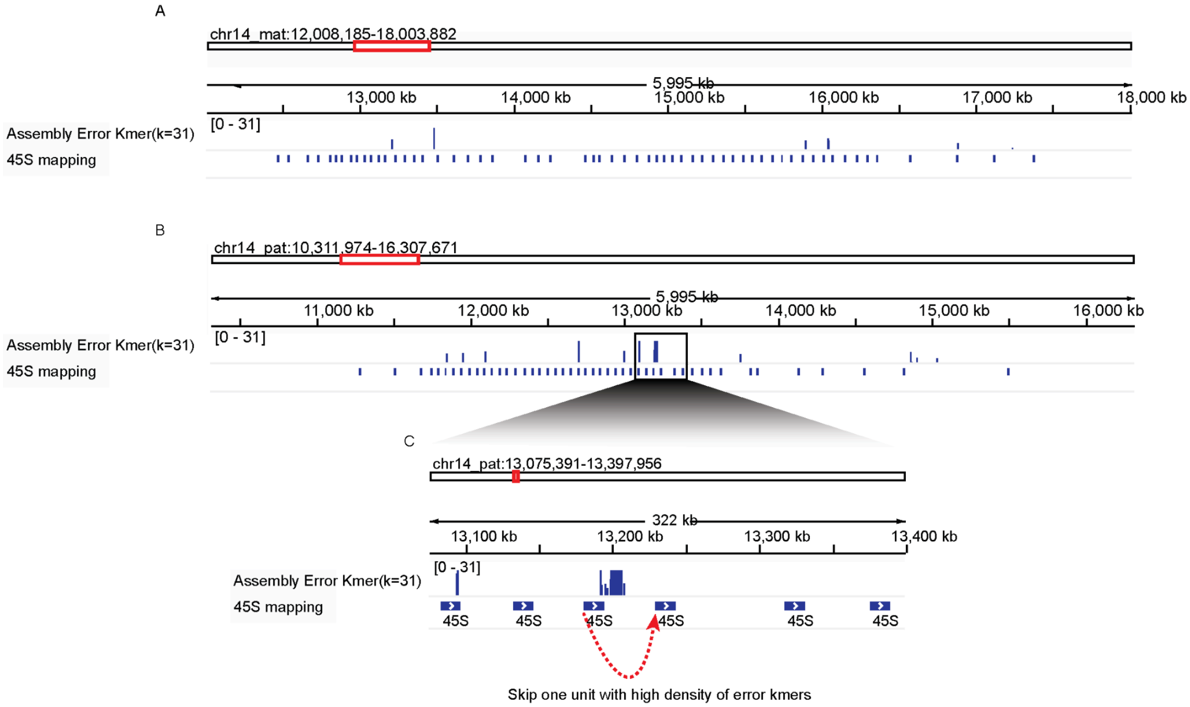


**Supplementary Figure 8. Error k-mer density in the rDNA region on chr14 after polishing.** A. rDNA (45S mapping) regions on the chr14 maternal haplotype. No regions show a high density of local error k-mers. B. rDNA regions on the chr14 paternal haplotype. One unit exhibits a high density of error k-mers. C. IGV zoom-in view of the boxed region in panel B. The red dotted arrow indicates the site where the assembly was rejoined after removal of the erroneous unit.

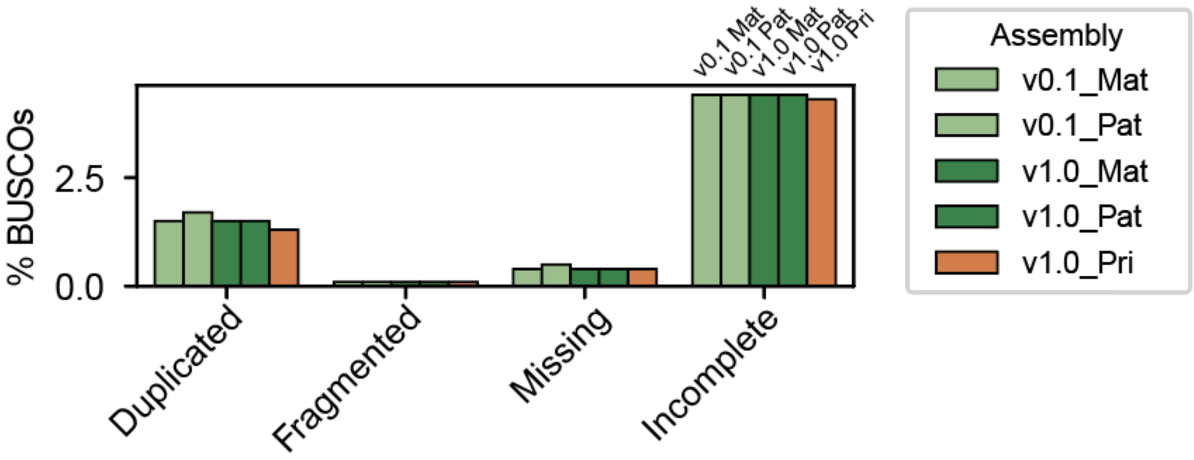


**Supplementary Figure 9. BUSCO scores percentages excluding the “Single Copy” category.**
This comparison shows BUSCO results using the cetartiodactyla_odb10 database, excluding the “Single Copy” category. “Incomplete” indicates predictions containing internal stop codons in Miniprot gene predictions.

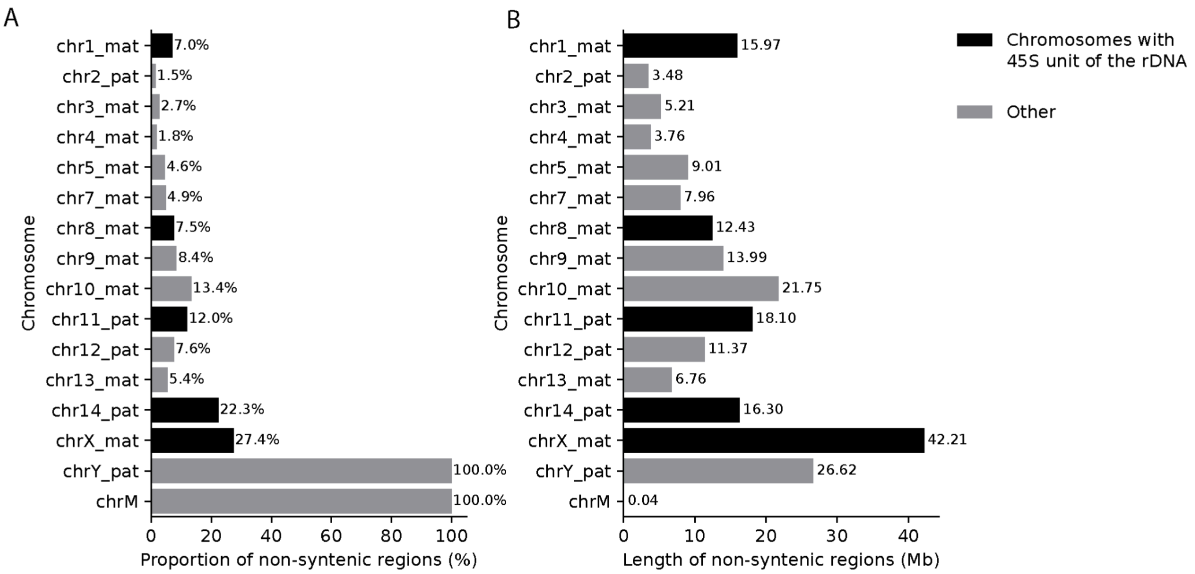


**Supplementary Figure 10. Length and proportion of non-syntenic regions for each chromosome in v1.0-Pri compared to ASM1759144v1.**

Non-syntenic regions were defined as unmappable regions when aligning T2T-mGirTip1v1.0 Primary assembly to ASM1759144v1.Chromosomes with >85% identity to the human 45S rDNA unit are highlighted in black. (A) Proportion of non-syntenic regions. (B) Total length of non-syntenic regions in Mbp.

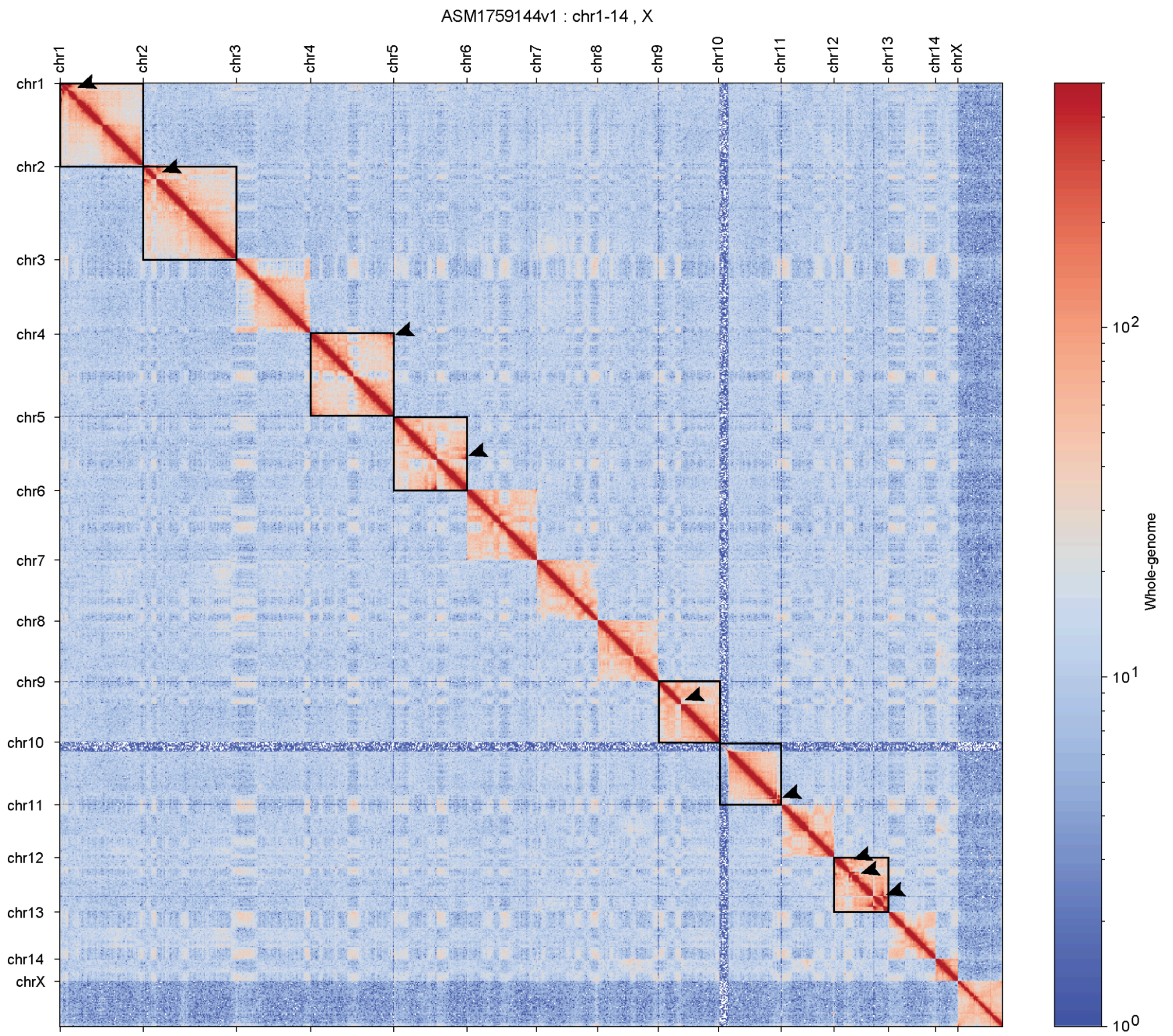


**Supplementary Figure 11. Whole-genome Hi-C contact map of mGirTip1 aligned to ASM1759144v1.**

The Hi-C contact map reveals large structural differences, including detectable inversions on chr1, 2, 4, 9, and 10. A more detailed zoom-in of these regions is provided in Supplementary Figure 14. Because ASM1759144v1 lacks chrY and chrM, only the 14 autosomes and chrX are shown here; other short unplaced scaffolds are not displayed. Black boxes highlight the chromosomes identified as having inversions in Figure 6, and arrows indicate long-range interactions within the chromosomes.

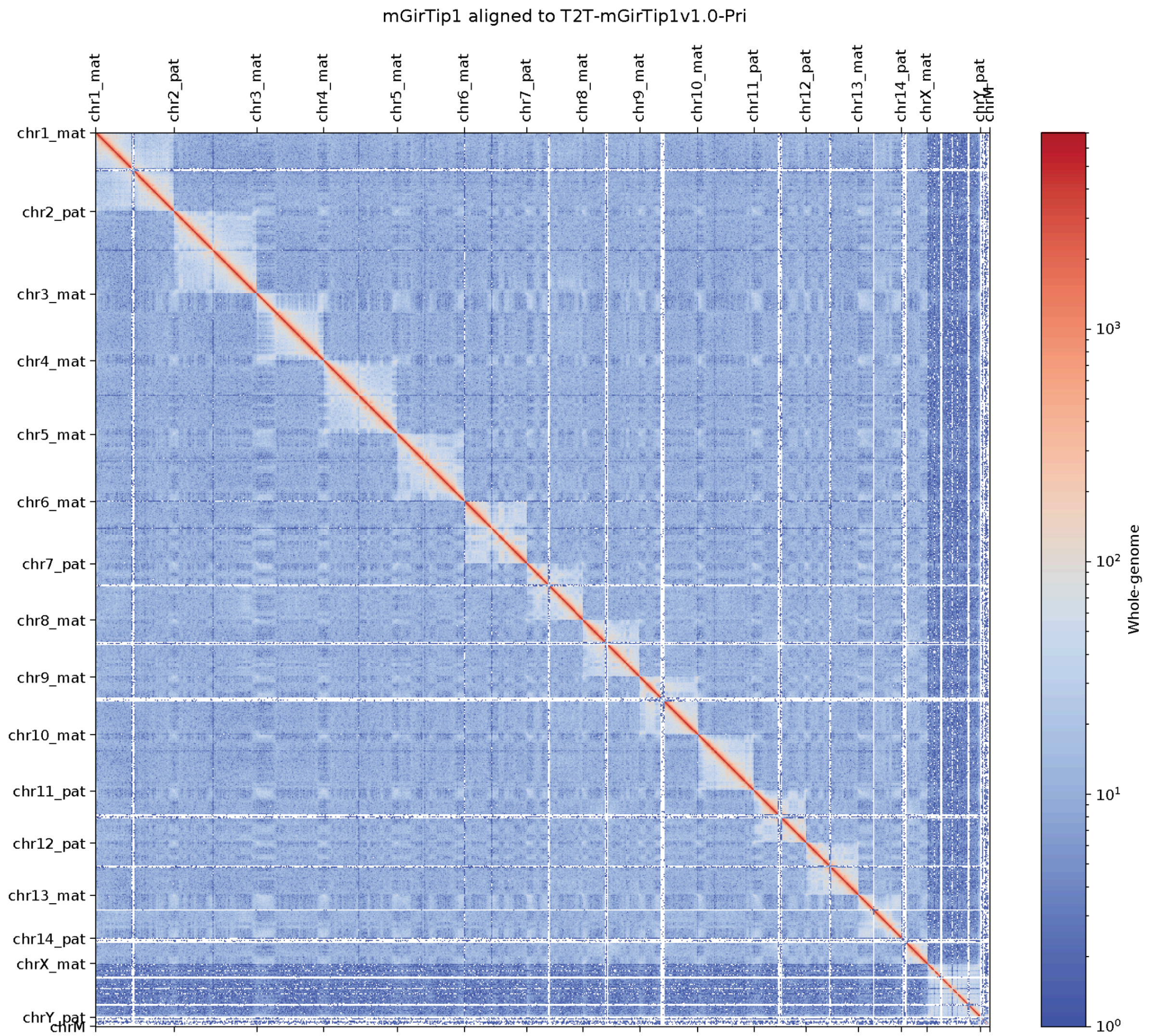


**Supplementary Figure 12. Whole-genome Hi-C contact map of mGirTip1 aligned to the T2T-mGirTip1v1.0 Primary assembly.**

The Hi-C contact map shows no structural variants across the whole genome. All 14 autosomes, the 2 sex chromosomes, and chrM are displayed.

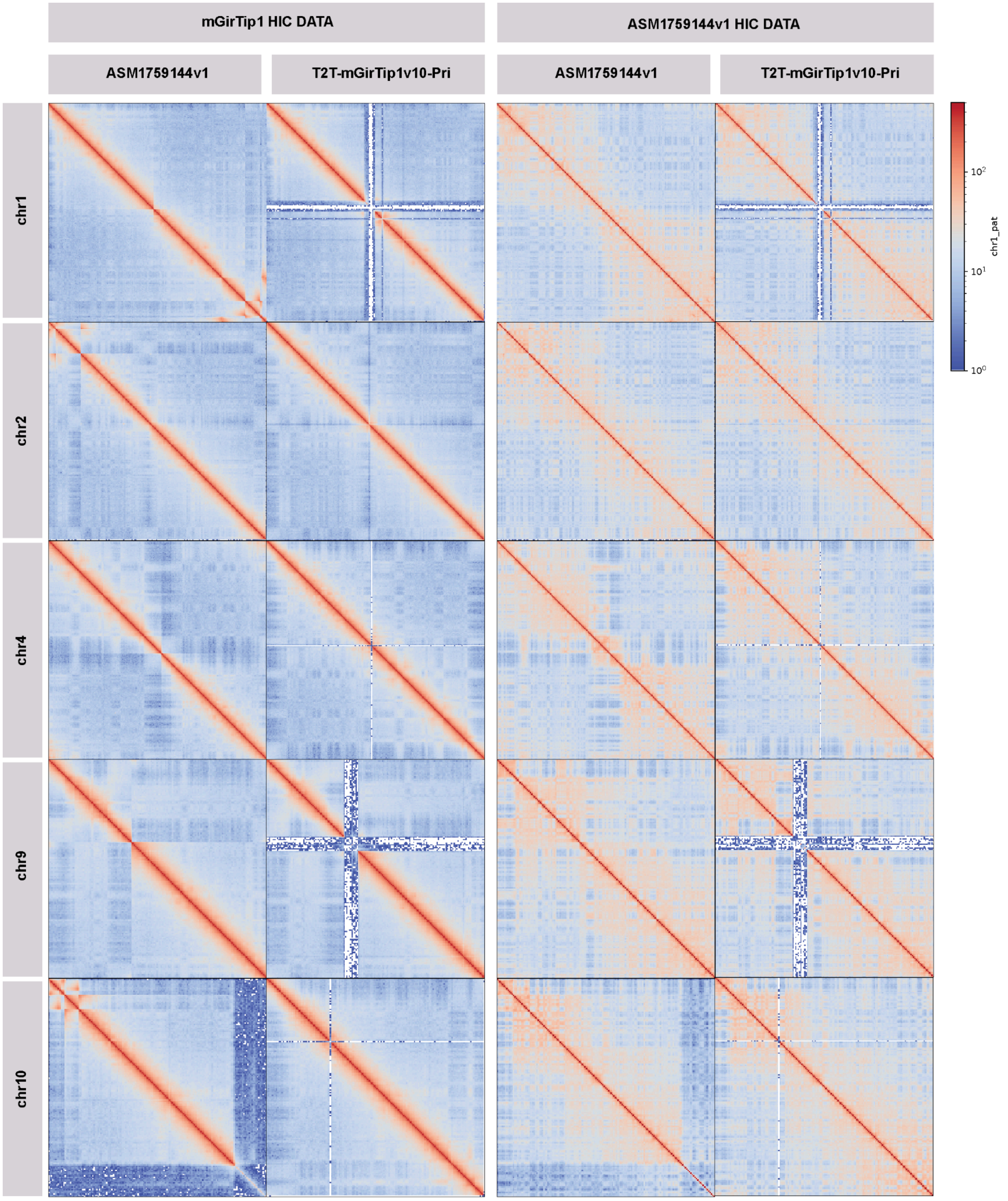


**Supplementary Figure 13. Large structural errors in the ASM1759144v1 assembly.**
Hi-C contact maps are shown for the full regions of each contig. Contigs with large structural differences between ASM1759144v1 and v1.0-Pri, as detected by mapping, are highlighted in each row (chr1, chr2, chr4, chr9, and chr10). Hi-C data from mGirTip1v1 is shown in the two columns on the left, while Hi-C data from ASM1759144v1 is shown in the two columns on the right. For each dataset, they are divided according to the reference used (ASM1759144v1 or v1.0-Pri). Hi-C data from ASM1759144v1 shows better concordance to v1.0-Pri despite having more long-range interactions, indicating the Hi-C scaffolding likely mis-oriented the sequences within ASM1759144v1.

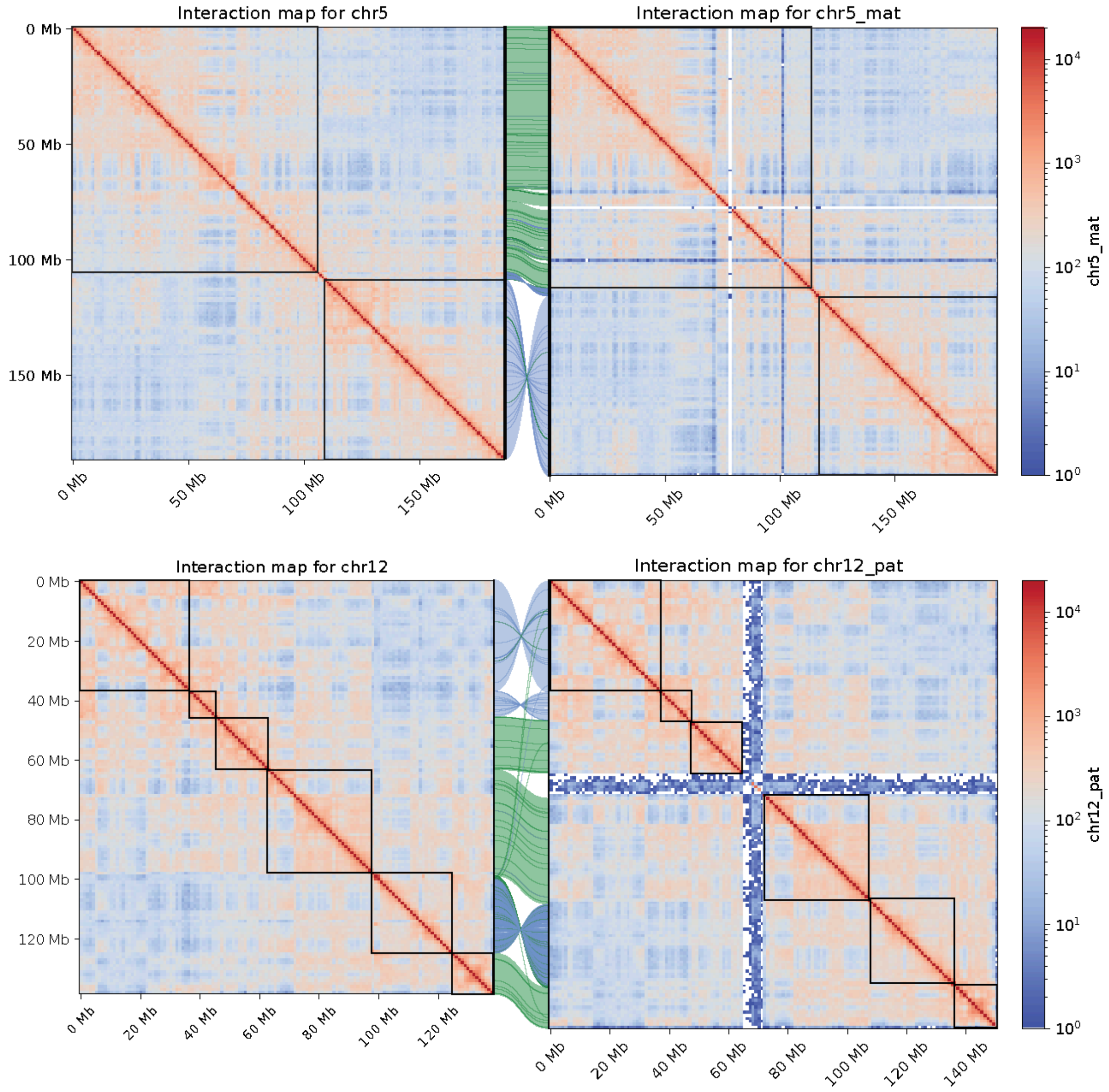


**Supplementary Figure 14. Hi-C contact map of ASM1759144v1 data for chr5 and chr12.** This figure corresponds to the same region shown in Figure 6G and 6H using Hi-C data from ASM1759144v1. Regions of large alignment blocks are highlighted with individual black boxes matching the corresponding alignments.
